# Unraveling the Molecular mechanism of Polysaccharide Lyases for Efficient Alginate Degradation

**DOI:** 10.1101/2024.08.02.606366

**Authors:** José Pablo Rivas-Fernández, Marlene Vuillemin, Bo Pilgaard, Leesa J. Klau, Folmer Fredslund, Charlotte Lund-Hanssen, Ditte H. Welner, Anne S. Meyer, J. Preben Morth, Flora Meilleur, Finn L. Aachmann, Carme Rovira, Casper Wilkens

## Abstract

Alginate lyases (ALs) are essential for breaking down brown macroalgae alginates, widely used naturally-occurring polysaccharides. Their molecular mechanisms remain challenging due to the lack of catalytically competent Michaelis-Menten complex structures. We here provide structural snap-shots and dissect the mechanism of mannuronan-specific ALs from family 7 polysaccharide lyases (PL7), employing time-resolved NMR, X-ray, neutron crystallography, and QM/MM simulations. We reveal the protonation state of critical active site residues, enabling atomic-level analysis of the reaction coordinate. Our approach reveals an endolytic and asynchronous *syn* β-elimination reaction, with Tyr serving as both Brønsted base and acid, involving a carbanion-type of transition state. This study not only reconciles previous structural and kinetic discrepancies, but also establishes a comprehensive PL reaction mechanism applicable across lyase families, which can guide the engineering of ALs for tailored alginate oligosaccharide production.

## Introduction

Alginates are linear anionic polysaccharides of 1,4-linked β-D-mannuronic acid (M) and its C-5-epimer α-L-guluronic acid (G) (Figure 1A)^1^. They are found in brown seaweed cells, where they account for 10–45% of the dry weight^2^, but can also be produced by bacteria from the genera *Azotobacter* and *Pseudomonas* as an exopolysaccharide^3^. Alginates are synthesized as polyM, and they can be modified by alginate C5-epimerases at the polymer level resulting in the formation of the different G- and mixed MG blocks. Such modifications greatly affect the physicochemical properties of the alginate polysaccharides, as the G blocks can bind Ca^2+^ (and other divalent ions) resulting in the formation of hydrogels^4^. Due to their physicochemical properties, alginates have attracted industrial interest, notably in food and biomedical applications^5–7^. Approximately 35,000 metric tons of alginates and alginate oligosaccharides are produced industrially per year by extraction from harvested or farmed brown seaweed, with a global market value of about US$450 million^8,9^.

**Figure 1.**
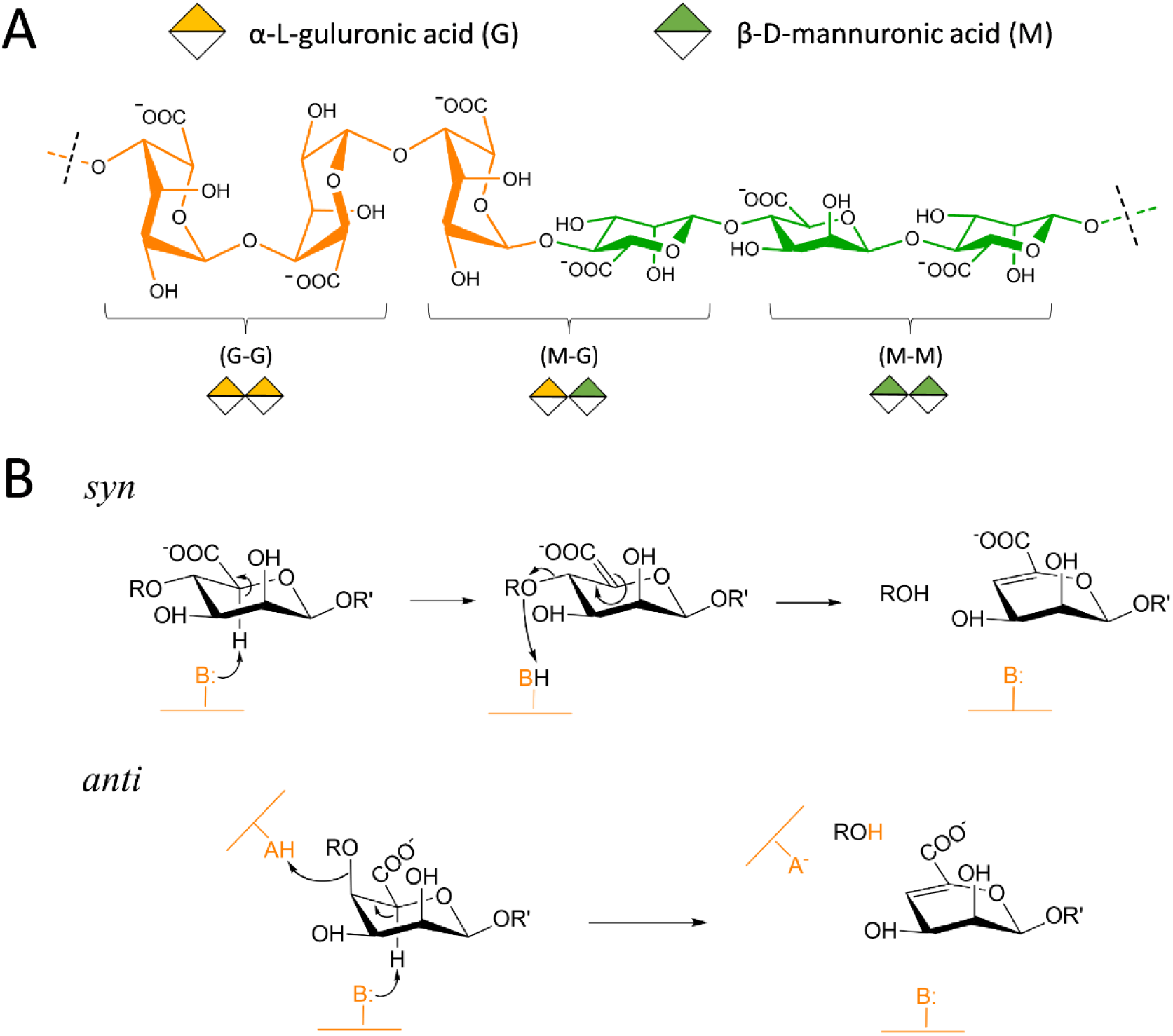
A) Chemical structure of alginates. The unit blocks, α-L-guluronic acid and β-D-mannuronic acid, are colored in orange and green respectively. B) Chemical reactions catalyzed by alginate lyases (Als). *Syn* reaction is presented above, while *anti* reaction is shown below. Acid and base residues, which belong to the enzyme, are denoted as A and B respectively and colored in light orange.

The degradation of alginates is catalyzed by alginate lyases (ALs), which are reported in 15 of the 41 polysaccharide lyase (PL) families in the Carbohydrate Active enZyme database (CAZy)^10^. These enzymes can be classified either as *endo*-acting, which cleave the alginate polysaccharides in the middle of the chain, initially releasing alginate oligosaccharides (AOS) that often are further degraded to di- and tri-AOS, or *exo*-acting, which cleave the polysaccharide, or more often the AOSs, by attacking the terminal end to liberate a monomer ^11–13^ (Figure 1B). The increased use of alginates in industrial applications have led to an interest in controlling their modification by enzymatic processing. However, our understanding of the mechanisms by which enzymes catalyze alginate modifications is incomplete due to lack of catalytically competent Michaelis-Menten structures in complex with substrate and products, limiting our ability to tailor-make alginates and AOSs^7,14,15^.

PLs cleave glycosidic bonds *via* an elimination reaction, described as a β-elimination reaction of the O’-C4 glycosidic bond, leading to formation of the 4,5-unsaturated sugar 4-deoxy-L-erythro-hex-4-enopyranosyluronic acid (denoted as Δ) at the non-reducing end of the released product^11,16^ (Figure 1B). A catalytic base (usually a negatively charged tyrosine or a histidine)^15^ is expected to abstract the H5 proton from the sugar moiety at subsite *+1*, resulting in the formation of an unsaturated bond between C4 and C5. The bond cleavage is assisted by the transfer of a proton from a general acid to the glycosidic oxygen^16^ (Figure 1B). The reaction can take place in *syn* or *anti*, depending on whether the abstracted proton is on the same side of the departing glycosidic oxygen (*syn*) or it is on the opposite side (*anti*), with respect to the sugar ring (Figure 1B). In the *syn*-elimination, a single residue, typically a tyrosine^17^, acts as both catalytic base and acid^17,18^, whereas two residues are needed when the reaction is *anti* ^3,15^. The identity of the catalytic base and the general acid remains unknown for several PLs. In addition, fundamental aspects of the catalysis such as whether the reaction is concerted or step-wise are unanswered.

The PL7 family holds the largest number of ALs that degrade alginates. The PL7’s harbors members that specifically degrade M, G or MG blocks in alginates, and some that are able to degrade two or three different blocks in *endo* or *exo* fashion^12,15,17^. Crystal structures, mainly originating from bacterial enzymes, have revealed that PL7 enzymes display a β-jelly roll fold (Figure 2A–D)^12,17^, which is shared with other PL families such as PL18^19^ and PL36^20^, that forms a groove harboring the active site (Figure 2A–F). Some PL7 members, such as the G-specific enzyme from *Sphingomonas Sp.* A1, have been suggested to employ an *anti* β-elimination mechanism^21,22^. Other PL7 enzymes, such as *Flavobacterium* sp. alginate lyase A (FlAlyA), were suggested to follow a *syn* β-elimination to degrade its preferred substrate polyM, based on crystal structure data and mutational analysis^17^. Such a mechanism was also recently suggested to apply to the β-D-glucuronan lyase TpPL7A from the fungus *Trichoderma parareesei*^23^. However, the reaction mechanism of ALs cannot be revealed due to either low resolution of the crystal structures or misplacement of substrates from subsites *-1* and *+1* due to enzyme mutations^21,22,24^. A recent high resolution^24^ crystal structure of a PyAly from the alga *Neopyropia yezoensis* in complex with PentaM revealed that two loops around the substrate binding cleft determine the distribution of products. Unfortunately, catalytically relevant substrate-enzyme interactions at the *-1* and *+1* subsites could not be analyzed due to mutation of critical active site residues (H125A/Y223A). Therefore, detailed characterization of the active site and the catalytic mechanism of PL7 enzymes and ALs in general remains elusive.

**Figure 2.**
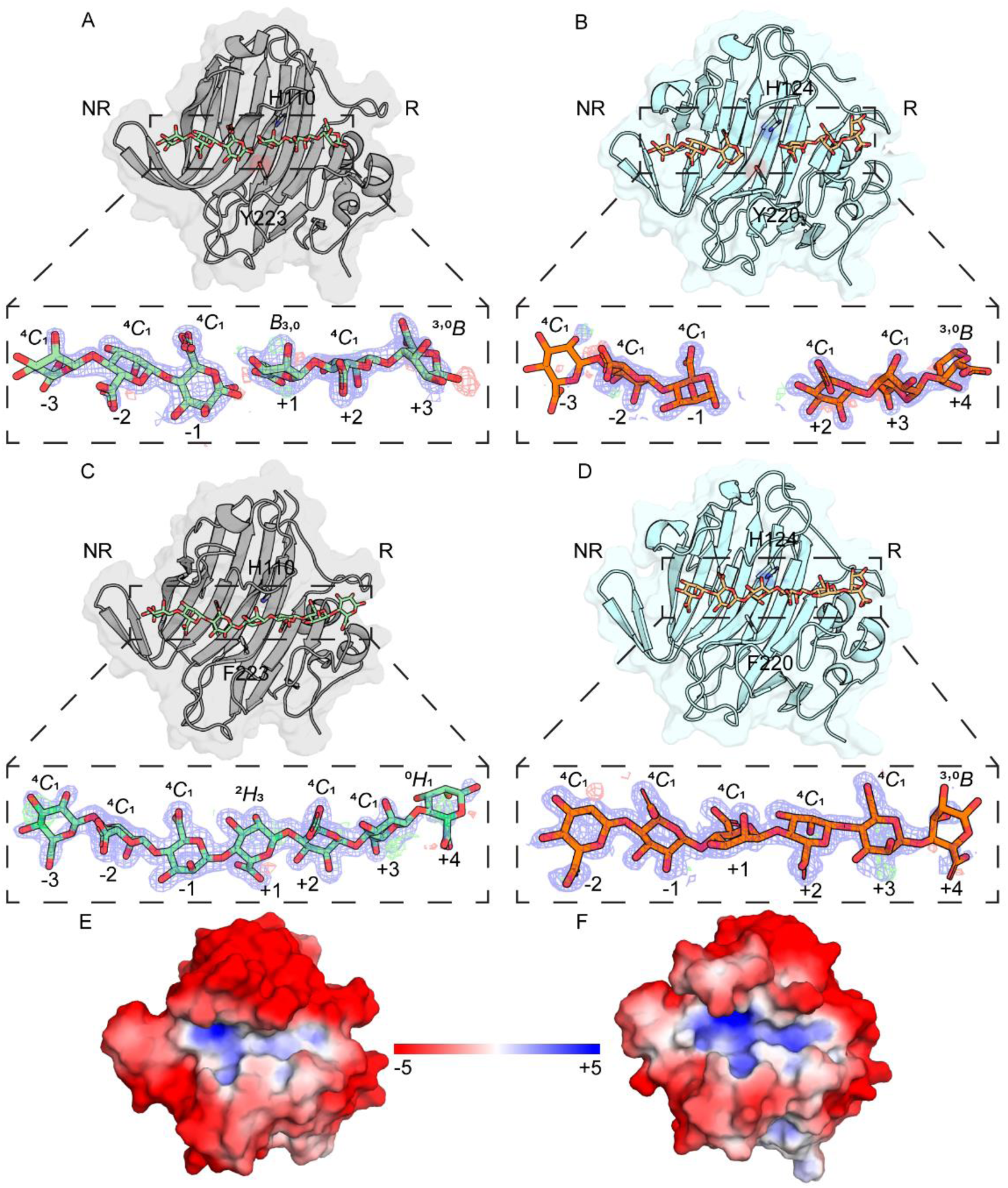
A) *Ps*Alg7A (grey) in complex with PentaM (green), B) *Ps*Alg7C (cyan) in complex with HexaM (orange), C) *Ps*Alg7A-Y223F (grey) in complex with HexaM (green) and D) *Ps*Alg7C-Y220F (cyan) in complex with HexaM (orange), E) electrostatic plot of *Ps*Alg7A at pH 5 and F) electrostatic plot of *Ps*Alg7C at pH 8. Electron density maps, 2fcfo (blue mesh) contoured to 1.0σ with a cutoff at 1.6 Å for C and fcfo (green and red) contoured to +3.0 and -3.0σ, respectively with a cutoff at 1.6 Å for C. NR and R indicates the non-reducing end and reducing end, respectively.

Here we employ a multidisciplinary approach including single crystals X-ray and neutron diffraction, NMR spectroscopy and QM/MM simulations to uncover substrate recognition and the molecular mechanism for M-specific PL7 enzymes. Time-resolved NMR was used to follow the reactions of two PL7 members *Ps*Alg7A and *Ps*Alg7C from the marine fungus *Paradendryphiella salina*, affirming that they are endolytic and M specific. In addition, we obtained high resolution X-ray crystal structures of the of the two alginate lyases in complex with M-oligosaccharides which, combined with neutron diffraction structures of *Ps*Alg7C, informed QM/MM simulations of *Ps*Alg7A that provided a precise atomistic picture of the reaction coordinate. The simulations confirm the *syn* β-elimination mechanism and reveal that the reaction takes place in one step, with a tyrosine acting as both catalytic base and acid, and involves the formation of a carbanion-type species at the transition state. Altogether, we provide an atomic description of the mechanism of action of a PL acting on alginate.

## Results

### Biochemical characterization of PsAlg7A and C

The endolytic enzymes *Ps*Alg7A and C are part of the marine fungus *P. salina* metabolization of alginates where *Ps*Alg7C (pH optimum 8) most likely takes part in the saprophyte’s initial attack on the seaweeds cell wall, while *Ps*Alg7A (pH optimum 5), together with an exolytic PL8 alginate lyase, finalize the degradation of the M blocks in alginates^13,25^.

The mode of action and product formation for *Ps*Alg7A was studied with time-resolved NMR on ^13^C-1 label poly-M substrate. *Ps*Alg7A produced oligomers with unsaturated end (Δ) as small as dimers, but no unsaturated monomer was formed (Figure 3 and Supplementary Figure 1). The signal for the unsaturated end (C-1Δ at 102.8 ppm) in trimers or oligomers (>DP 2) was present from the beginning of the reaction, while a slowly increasing signal for the unsaturated end (C-1Δ at 102.5 ppm) in dimers appeared later. Based on the integrals (for C-1 of unsaturated ends and ratio conformation), the distribution of the reaction products are ∼84% trimers or oligomers and ∼16% dimer. Looking at the formation of reducing ends, the signals α (96.2 ppm) and β (96.3 ppm) conformers in the unsaturated trimers or oligomerswere present from the start of the reaction, while the α reducing end (95.9 ppm) of dimers appeared later. Altogether, the time-resolved NMR results strongly indicate that *Ps*Alg7A displays an endolytic mode of action, producing mainly unsaturated trimers or oligomers with a higher degree of oligomerization on the polyM substrate. *Ps*Alg7C exhibits a similar endolytic mode of action and product formation (Supplementary Figures 2 and 3).

**Figure 3.**
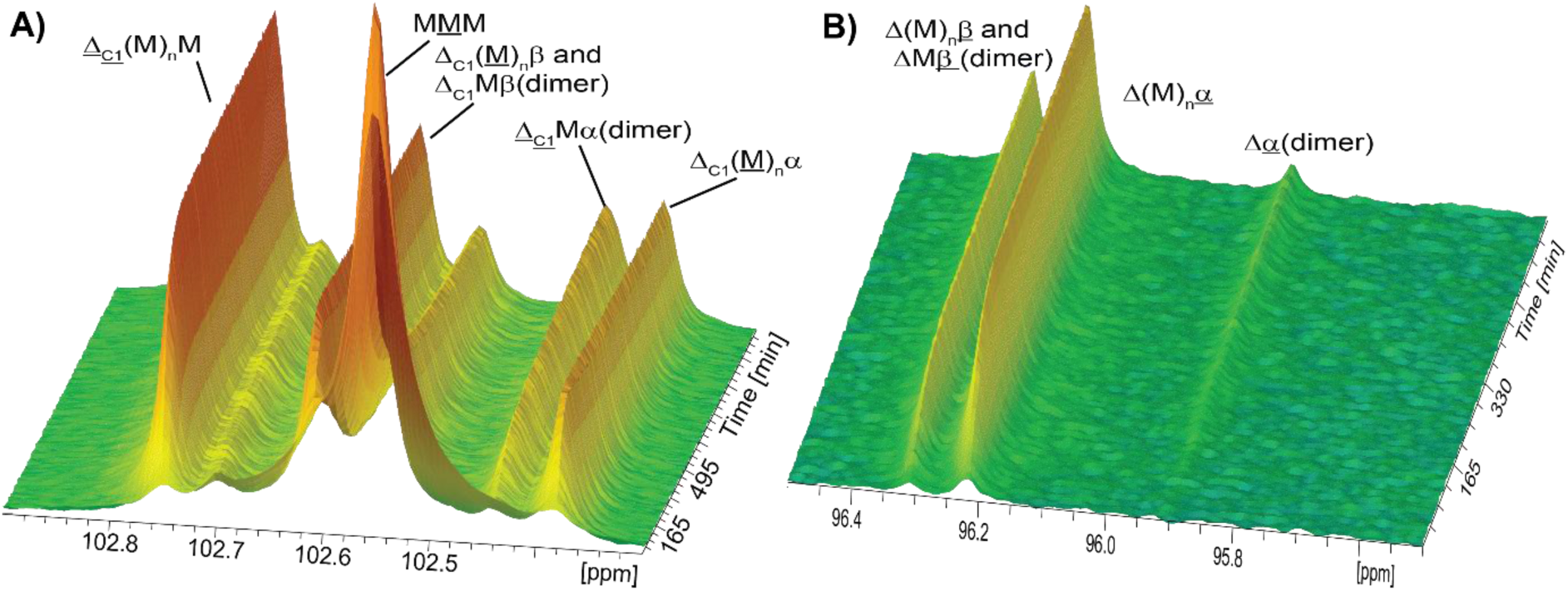
Time-resolved ^13^C-NMR spectra of ^13^C-1-enriched polyM treated with *Ps*PL7A. Reaction was conducted in 3mm NMR tube with 160 μL of substrate buffer solution (15 mg/mL ^13^C-1-polyM substrate in 5 mM Na-acetate, pD 5.5 with 100 mM NaCl and 1.5 mM ZnCl_2_ in 99.9% D_2_O) and 10 µL of 147.5 mg/mL *Ps*PL7A incubated at 25 °C. Pseudo 2D time-resolved spectra recorded by acquiring 1D carbon spectra every 5 mins for a total of 21 hrs 20 mins. Panel A) shows anomeric region and B) reducing end region in the carbon spectra. α and β indicate the signals for the reducing end of alginate, M is mannuronate, C1 indicate anomeric in alginate sugar ring, Δ: 4,5-unsaturated 4-deoxy-L-*erythro*-hex-4-enepyranosyluronate, underlined indicate the residue with the anomeric C1 giving rise to the signal.

### Overall structure of PsAlg7A and C and their relation to other PL7 members

The crystal structures of *Ps*Alg7A and C were determined at 0.82–2.00 Å resolution (data processing statistics are summarized in Supplementary Table 1). Overall, the electron density maps are for the proteins and the ligands was found to be well defined (Figures 2A–B, E–F). The tertiary structure of *Ps*Alg7A and C resemble that of other PL7 members, which are composed of single β-jelly roll domain with an inner and outer convex anti-parallel β-sheet (Figures 2A–D). Structural analysis of the Protein Data Bank (www.pdb.org)^26^ using the DALI server^27^ revealed that the closest structural homolog for *Ps*Alg7A is indeed *Ps*Alg7C (C_α_ r.m.s.d. is 0.37 Å, for 166 atom pairs). *Ps*Alg7A and C might be members of a yet to be defined PL7 subfamily in CAZy^13^, as they exhibit distinct structural features from other PL7 members. The structure of *Ps*Alg7A and C differ from other structurally determined PL7 members, except for PyAly from algae *Neopyropia yezoensis* (PDB ID 7W13)^24^, by the presence of two extended β-strings at the non-reducing end of the binding cleft (*Ps*Alg7A: V142–I147 and V150–V155. *Ps*Alg7C: I136–L141 and V144–F149) that form a lid-like structure on top of the substrate (Figures 2A–D). Attempts to unravel the importance of these extensions was hindered by the lack of soluble protein during expression, which precluded us from drawing any conclusions regarding their influence on the enzyme activity (data not shown). However, the crystal structures show that the two β-strings form interactions with the HexaM substrate in *Ps*Alg7A (Figures 2A and C), implying that they are probably involved in substrate binding. These two additional β-strings are not present in the β-glucuronan lyase *Tp*PL7A from the terrestrial fungus *T. parareesei*^23^, suggesting that this extension of the active site grove could be limited to alginate lyases from marine eukaryotes or may define substrate selectivity.

### Substrate and product accommodation

The substrate binding groove of ALs is in general positively charged, which favors the binding of the negatively charged substrate^15^. This is also the case for *Ps*Alg7C (Figure 2F). However, the binding groove is less positively charged for *Ps*Alg7A (Figure 2E), which may explain the seven-fold higher *K*_m_ of *Ps*Alg7A (17.4 mM) compared to *Ps*Alg7C (2.4 mM) towards polyM^13^. This could also explain the large difference in catalytic turnover rates, with *Ps*Alg7C acting more than three times faster on polyM than *Ps*Alg7A.

Structures of five PL7 AL in complex with different alginate oligosaccharides have been reported, revealing a hydrogen bond network and stacking-like interactions important for positioning the substrate in the active site groove^21,22,24^. All these complex structures were obtained for catalytically inactive mutants, in which one or more putative essential residues were mutated. Here, we were able to soak M oligosaccharides with a degree of polymerization (DP) 4–6 into crystals of *Ps*Alg7A, resulting in *in crystallo* cleavage of the sugars (Figure 2A) (1.46–1.87 Å resolution, Supplementary Table 1). This confirmed the *-1* and *+1* subsites for PL7 family members that define the location of the scissile bond (Figure 2A). Previously, these subsites were assumed based on the positions of the presumed catalytic tyrosine and histidine relative to the uronic acids in the complex structures^21,22,24^. The negative subsites (Figure 2A) indicate the position of the non-reducing end of the substrate, while the positive subsites indicate the reducing end^28^. TetraM, PentaM and HexaM oligosaccharides were found to span subsites *-3* to *+3*, suggesting that TetraM and PentaM both are accommodated in two or more different ways in the active site.

Attempts to perform *in crystallo* substrate cleavage for *Ps*Alg7C were unsuccessful. The structures, determined at 1.09–1.24 Å resolution, showed that M oligosaccharides (DP 2–6) are present at negative and positive subsites, but the *+1* subsite is unoccupied (Table S2; Figure 2B). However, this subsite is occupied in both inactive mutants *Ps*Alg7C-H124N and *Ps*Alg7C-Y220F, which crystallized in space groups *C*12_1_ and *P*12_1_1, and *P*1, respectively (Supplementary Table 1), where, as *Ps*Alg7C, crystallized in space group *P*2_1_2_1_2_1_. A loop from the neighboring molecule in the crystal lattice containing S228 was found to pose a steric hindrance for the carboxylic group of the M moiety at subsite +*1*, precluding productive binding. A G moiety was consistently observed at subsite +*3* (Figure 2B), which may stem from impurities in M oligosaccharides, suggesting that *Ps*Alg7C has a preference for G moieties at this subsite. The G moiety was also observed at subsite +*3* in *Ps*Alg7C-H124N in complex with PentaM.

To obtain Michaelis-Menten-like complexes of *Ps*Alg7A and *Ps*Alg7C, the inactive mutants *Ps*Alg7A-Y223F, *Ps*Alg7C-H124N and *Ps*Alg7C-Y220F in complex with M oligosaccharides (TetraM and PentaM) were crystallized. The corresponding structures were determined at 1.45–1.81 Å (*Ps*Alg7A-Y223F), 1.05–1.51 Å (*Ps*Alg7C-Y220F) and 0.87–1.13 Å resolution (*Ps*Alg7C-H124N). The structures show that the TetraM and PentaM substrates span subsites *-2* to *+3* in *Ps*Alg7A-Y223F (Supplementary Table 2), while HexaM spans *-3* to *+4* (Figure 2C; Supplementary Table 2). As found for the product complexes, this suggests that the M oligosaccharides can be accommodated in two or more different ways in the active site. Except for G moieties observed at subsite -*2*, the M moieties in the *Ps*Alg7C complex structures resemble those observed in the *Ps*Alg7C-Y220F structure in complex with HexaM (Figure 2D).

The structure of *Ps*Alg7A-Y223F in complex with HexaM reveals a highly complex hydrogen bond network, which is probably responsible for the substrate accommodation in the active site groove (Supplementary Figure 4), optimizing enzyme interactions with the M moieties. The majority of the enzyme-substrate interactions are mediated by water molecules, while some direct hydrogen bonds are observed (Supplementary Figure 4). A similar hydrogen bonding network was observed in the structures of *Ps*Alg7A-Y223F in complex with TetraM and PentaM, respectively, confirming its importance for substrate recognition.

Most of the M moieties of the *Ps*Alg7A-Y223F-HexaM complex are present in the ^4^*C*_1_ conformation, except the one at subsite *+1*, which adopts a distorted ^2^*H*_3_ conformation (Figure 2C). This is also the case on the Penta- and TetraM complexes (Supplementary Table 2). This unusual conformation^29^ is probably due to the Y223F mutation (subsequent QM/MM simulations on the wild-type enzyme, discussed below, supports this hypothesis).

In contrast to *Ps*Alg7A-Y223F-HexaM, all the β-D-mannuronic acid moieties are present in the ^4^*C*_1_ conformation in the *Ps*Alg7C-Y220F-HexaM complex (Figure 2D). However, the structure of *Ps*Alg7C-H124N shows that the mutation apparently results in a distortion of the M moieties (Figure 4A; Supplementary Table 2), which likely is a result of the loss of the interaction between H124 and the 2-OH group of the sugar unit at the *+1* subsite (Figure 4A), leading to a displacement of the scissile bond. In fact, N124 is observed in two different conformations when in complex with M oligosaccharides (Figure 4A), suggesting that the asparagine residue does not interact with the M moieties. This is likely the reason for the almost complete loss of activity upon mutating the wild-type histidine that was previously observed in other PL7 members^22,23^.

**Figure 4.**
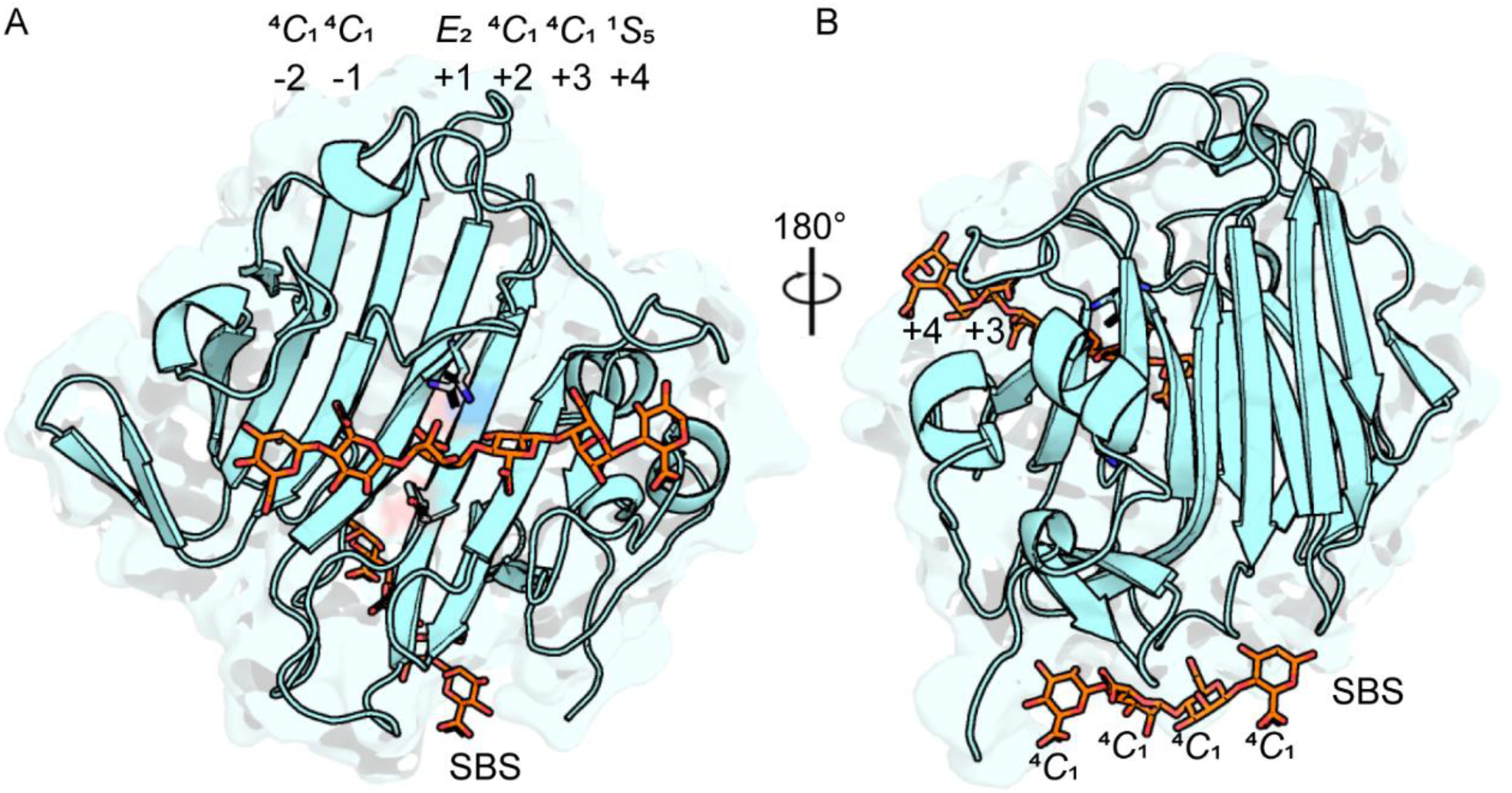
*Ps*Alg7C-H124N (cyan) soaked with PentaM (orange). A) Active site groove and B) the surface binding site (SBS). N124 and Y220 are shown as sticks.

### Surface binding site

A surface (or secondary) binding site (SBS) was observed in the crystal structure of *Ps*Alg7C-H124N in complex with PentaM on the opposite of the active site groove (Figure 4B). SBSs hold a diversity of functions and share many roles with the more known carbohydrate binding domains (CBMs), which are domains separate from the catalytic domains. SBSs are located at a certain distance of the active site on the catalytic domains, and some have previously been shown to be important for the activity and functionality of their parent catalytic domain ^30–32^, but no SBSs have been previously identified in PLs. Four M moieties are observed at the SBS on *Ps*Alg7C-H124N, which all interacts *Ps*Alg7C through hydrogen bonds (Figure 4B) that potentially can discriminate for M moieties. However, its role, if any, in *Ps*Alg7C’s degradation of M-blocks in alginate requires further investigations, which is beyond the scope of this study.

### Pre-organized active site

The overall structures of *Ps*Alg7A-Y223F-HexaM and *Ps*Alg7C-Y220F-HexaM complexes are rather similar. Their most noteworthy difference is R161 (*Ps*Alg7A) and R155 (*Ps*Alg7C) at subsites -*1* and +*1,* which are observed in different conformations (Supplementary Table 2 and Supplementary Figure 5). Another difference is that A169 (*Ps*Alg7A) at subsites -*2* and -*3* is replaced by the bulky F165 in *Ps*Alg7C. However, this does not affect the structure of the active site (Supplementary Figure 5), which displays similar hydrogen bonding interactions in both enzymes. Several water molecules mediating the hydrogen bonds between the M moieties at subsites -*1* and +*1* are observed in similar positions in both *Ps*Alg7A and C complexes (Supplementary Figure 5). This suggests that the active sites are preorganized for substrate binding and catalysis. Remarkably, replacement of F223 in *Ps*Alg7A-Y223F-HexaM by tyrosinate shows a clear short distance between the Tyr oxygen atom and the H5 atom of the saccharide at the *+1* subsite, supporting a role as general base in catalysis.

### Protonation states of putative catalytic residues

To investigate the protonation states of the residues interacting with the substrate, neutron diffraction structures of *Ps*Alg7C before and after soaking with PentaM were obtained. These structures were determined at 2.15 Å resolution (see Supplementary Table 3 for statistics). Of particular interest, the neutron structures revealed that Y220 is deprotonated (*i.e.* negatively charged) prior to substrate binding, but it is in its neutral form in the product complexes (deuterium occupancy was 0.73) (Figure 5B). Conversely, H124 holds only one proton (*i.e.* it is a neutral residue) in both the substrate-free structure and the product complex (Figure 5A– B). This strongly suggests that Y220 is the catalytic base that abstracts the proton from the saccharide at the *+1* subsite during catalysis.

**Figure 5.**
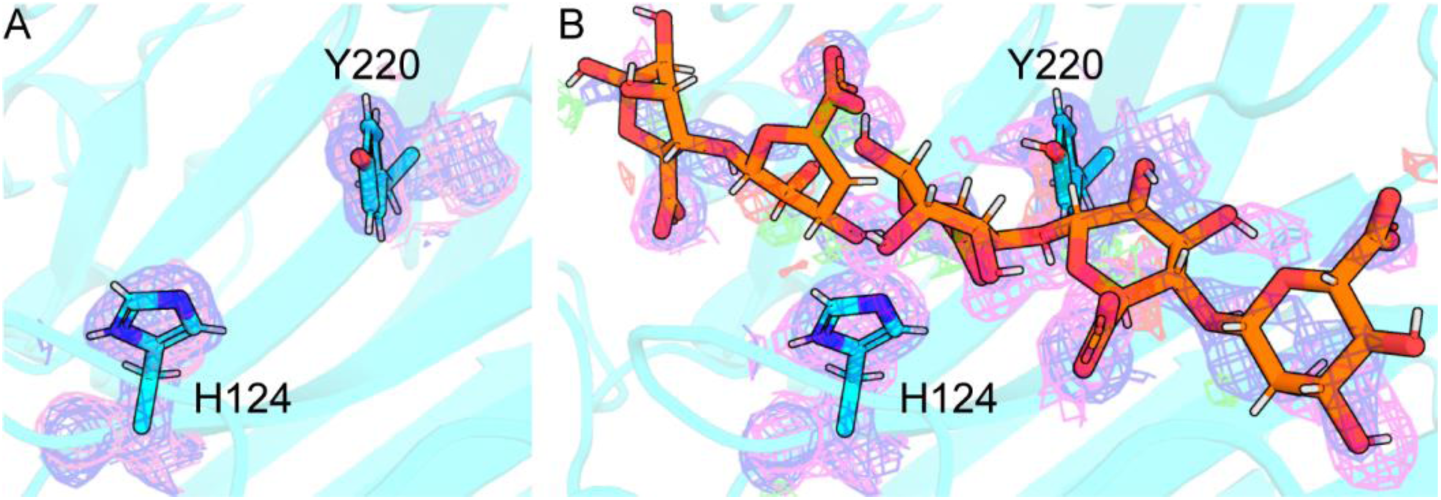
Protonation states of putative catalytic residues. Structure of A) *Ps*Alg7C and B) *Ps*Alg7C soaked with PentaM. Electron density maps from X-ray data, 2fcfo (blue mesh) contoured to 1.0σ with a cutoff at 1.6 Å for C and fcfo (green and red) contoured to +3.0 and -3.0σ, respectively with a cutoff at 1.6 Å for C, while electron density maps from neutron data, 2fcfo (magenta mesh) contoured to 1.0σ with a cutoff at 1.6 Å for C.

While in the neutral form, H124 cannot act as a proton donor, thus it cannot be the catalytic acid. This is consistent with the *syn*-elimination mechanism (Figure 1B), in which a tyrosine is expected to be the only catalytic residue. The present high resolution X-ray crystal structures indicate that the active site histidine (H124 in *Ps*Alg7C and H129 in *Ps*Alg7A) forms a hydrogen bond with the 2-OH of the M saccharide at subsite *+1* (Supplementary Figure 5), suggesting that it is in the N_δ_ tautomeric form. Together with the results obtained for *Ps*Alg7C-H124N, our results suggest that H124 is essential for a productive binding of the substrate but does not participate in the catalytic reaction.

### Partially protonated catalytic tyrosine in product complexes

The structures of *Ps*Alg7A and C in complex with DiM and TriM oligosaccharides, as well as those of enzyme mutants, show that the substrate spans only subsites *-3*, *-2* and -*1,* except for *Ps*Alg7C-Y220F in complex with TriM (Supplementary Table 2). This may explain why a build up of DiM and TriM is observed by time-resolved NMR spectroscopy (Figure 3 and Supplementary Figure 1). We therefore speculate that these structures mimic the enzyme state after leaving group departure, prior to a processive movement of the substrate in the active site groove. Similar to what was observed for *Ps*Alg7A in complex with the TetraM, PentaM and HexaM products (Figure 2A), the M moieties of DiM and TriM at subsite *-1* are often found in both α and β conformations (Supplementary Table 2). Y223 (*Ps*Alg7A) and Y220 (*Ps*Alg7C) are both within hydrogen bonding distance from the O1 atom of the M sugar at subsite -*1*. The observed α and β conformations may reflect that tyrosines Tyr223 and Tyr220 protonated, as found in the neutron structure of *Ps*Alg7C (Figure 5B).

### Conformation of the substrate β-mannurosyl unit at the +1 subsite

The structure of *Ps*Alg7A-Y223F in complex with HexaM (PDB ID 7NCZ) was used as initial template for the molecular dynamics (MD) simulations. The F223 residue was substituted for a tyrosine residue to reconstruct the native enzyme. The solution structure Michaelis-Menten complex (MC) was build using Amber 2020 tools (see details in the Methods section). The complex was subsequently minimized and equilibrated for 750 ns (Supplementary Figure 6). The simulations show that the enzyme-substrate complex remains stable in a catalytically productive configuration, with H129 and Y223 sandwiching the *+1* saccharide, which adopts a ^4^*C*_1_ conformation. The negatively charged oxygen of the putative catalytic base, Y223, is close to R82, forming a salt-bridge interaction. The O atom of Y223 is also very close to the H atom at C5, ready for proton abstraction.

To further confirm the conformation of the *+1* saccharide, we computed its conformational free energy landscape (FEL) using QM/MM metadynamics^33^. This approach has been previously used with success to elucidate substrate conformations in other CAZymes^34,35^. The use of a QM description of the substrate is necessary to exclude any bias due to the classical force-field, which often favors chair conformations^34^. The conformational FEL of the *+1* subsite M moiety in *Ps*Alg7A (Supplementary Figure 7) confirmed that ^4^*C*_1_ is the most stable conformation (the global minimum of the FEL). Remarkably, there is a particularly low-energy region that extends towards ^2^*S*_O_, which is precisely the conformation of the transition state (TS) of the reaction, as described below. Therefore, as previously observed for glycosidases^34^, the active site of *Ps*Alg7A is optimized to precisely stabilize the substrate in conformations that are relevant for catalysis.

### Reaction mechanism

One representative snapshot from the classical MD simulations was selected (Supplementary Figure 6) as starting point to initiate QM/MM MD simulations of the reaction mechanism. A large QM region (112 atoms) was taken as to include part of the substrate and the side chains of the main residues interacting with the *+1* saccharide (Y223, R82, K219 and Q127) (Supplementary Figure 8). The simulations were performed with the CP2K program^36^, using Density Functional Theory (DFT) at the QM region and the Amber-based force-fields at the MM region (see more details in the Methods section). The system was equilibrated for 7 ps at QM/MM MD level before starting the simulation of the reaction mechanism.

The chemical reaction was modeled using the OPES enhanced sampling method^37,38^, which provides a very efficient exploration of complex energy landscapes, as shown in recent applications to chemical reactions^39^. Two collective variables (CVs) were used to drive the system from reactants to products (see Figure 6A for CVs definition). The first CV was taken as the distance (d) of the scissile glycosidic bond [CV1 = d(O4-C4), where O4 refers to the O’ atom]. The second CV was taken as a combination of the distances involved in the possible motion of the catalytic proton (H5) [CV2 = d(C5-H5) – d(H5-O_Tyr223_) – d(O4-H5)]. Therefore, CV1 accounts for the cleavage of the glycosidic bond, while CV2 accounts for proton transfer events. Using these CVs, the QM/MM OPES simulation successfully drove the reactants towards the products of the reaction, *i.e.*, TriM and unsaturated (ManA)_4_. The computed reaction free energy landscape (Figure 6B) shows two well defined reactants and products minima, with the latter being lower in energy, separated by a unique transition state (TS). Therefore, the reaction is concerted and exothermic. The computed reaction free energy barrier for *Ps*Alg7A (16.5 kcal mol^-1^) is in very good agreement with the experimental value estimated from the reaction rate (17.3 kcal mol^-1^), which reinforces the mechanism obtained.

**Figure 6.**
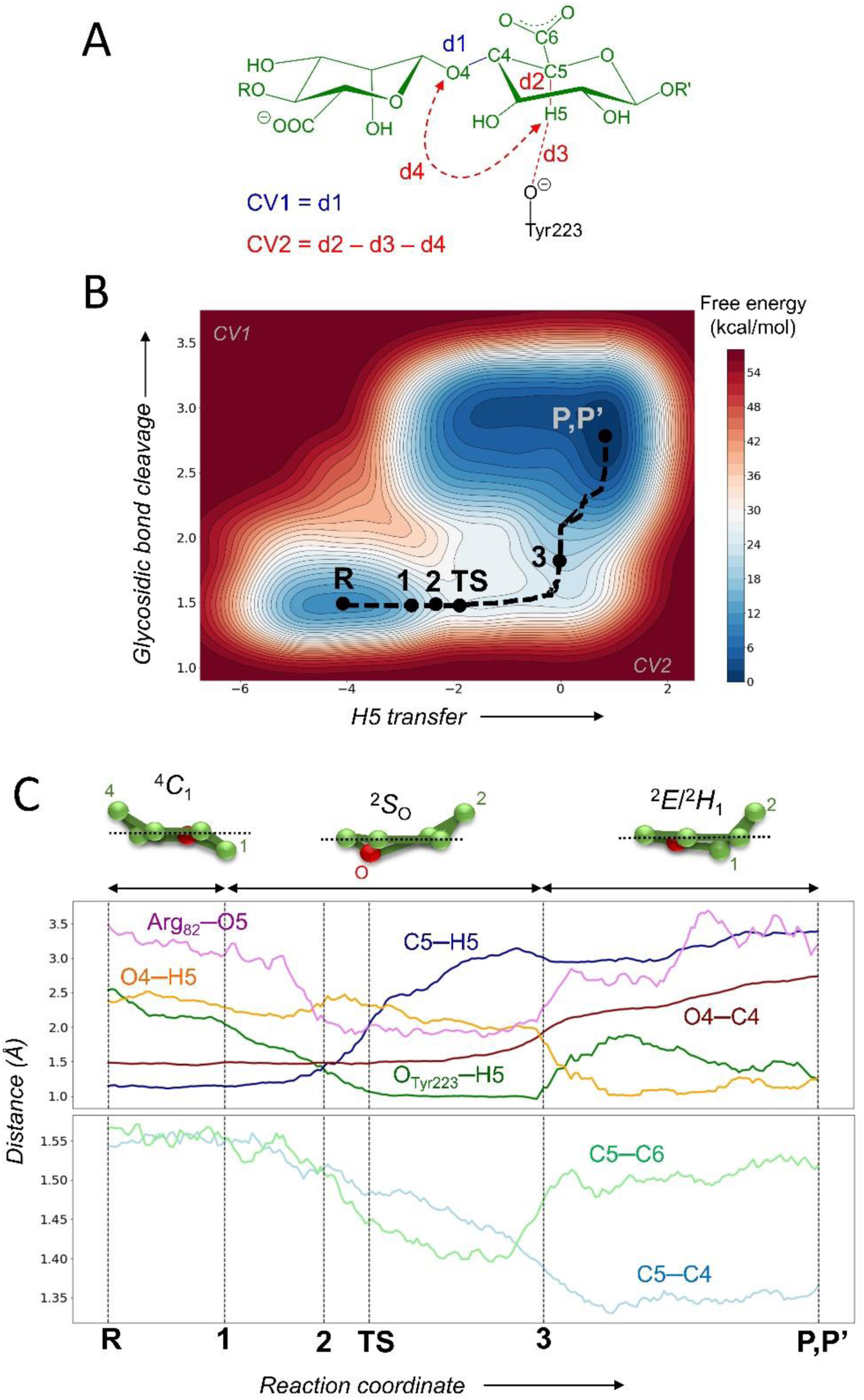
A) Collective variables used in the QM/MM OPES simulations of the catalytic mechanism of *Ps*Alg7A. B) Free energy landscape obtained from the simulations and minimum free energy pathway defining the reaction coordinate. C) Evolution of the most relevant distances along the reaction coordinate.

Analysis of the molecular changes along the minimum energy pathway (Figure 6C) shows that the reaction begins by a slight repositioning of R82 (state **1** in Figure 7), which detaches from Y223 and approaches the endocyclic O5 atom of the *+1* saccharide. This motion, although small, induces a distortion of the *+1* saccharide from the relaxed ^4^*C*_1_ chair to a skew-boat ^2^*S*_O_ conformation, enabling a close contact between the oxygen atom of Y223 and the H5 atom of the *+1* saccharide, which becomes ready for proton transfer. Subsequently, Y223 abstracts the H5 proton (state **2** in Figure 7) and the C5 atom acquires negative charge. The negatively charge on C5 is delocalized over C6, causing both the C4-C5 and C5-C6 bonds to shorten upon reaching the reaction TS (Figure 6C), which acquires carbanion character. The cleavage of the glycosidic O4-C4 bond takes place after the TS (state **3** in Figure 7), just before reaching the products state in which the glycosidic bond is fully broken. The *+1* sugar adopts an envelope (^2^*E*) conformation at the reaction products, due to the formation of a double bond between atoms C4 and C5 upon formation of the 4,5-unsaturated sugar 4-deoxy-L-erythro-hex-4-enopyranosyluronic acid. Therefore, even though the reaction free energy landscape (of Figure 6B) shows a unique TS, indicative of a concerted reaction, the reaction is highly asynchronous, with glycosidic bond cleavage taking place after proton transfer. Our simulations also predict a ^4^*C*_1_ → [^2^*S*_O_]^‡^ → ^2^*E* conformational itinerary for the β-elimination reaction catalysed by *Ps*Alg7A.

**Figure 7.**
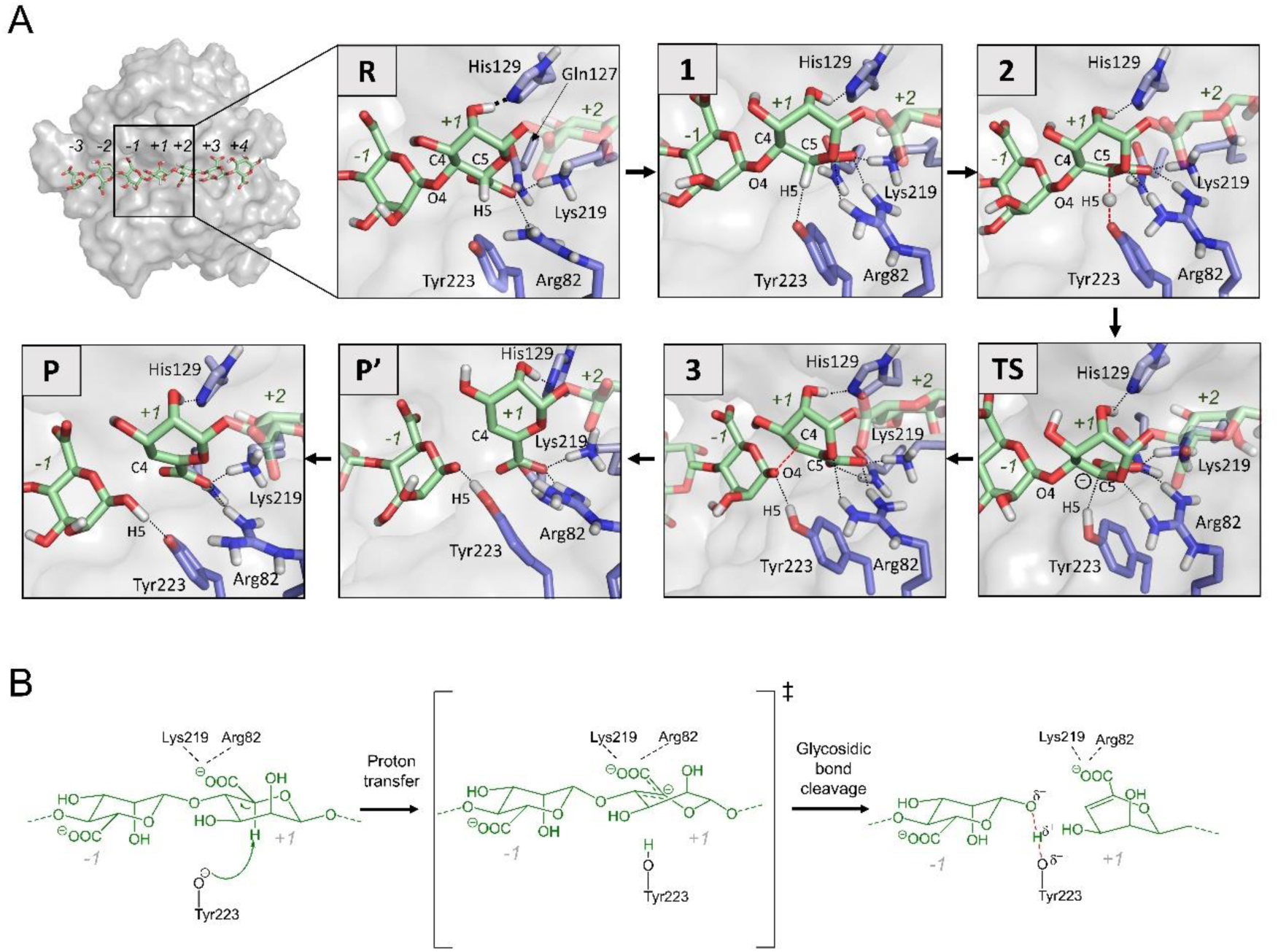
A) Representative structures along the reaction coordinate, obtained from QM/MM OPES simulations. Hydrogen atoms that are attached to C atoms have been omitted for clarity, except the H5 atom. (B) Proposed reaction mechanism for *Ps*Alg7A according to all results of this work

Further analysis of the species present at the products state (P) reveals that it is a mixture of states in which Y223 can be either neutral or deprotonated. In other words, the H5 proton can be either located at the oxygen atom of Y223 (with the new reducing end being an alkoxide) or at the oxygen atom of the new reducing end (with Y223 being negatively charged). A separate QM/MM MD simulation of the reaction products confirmed that the proton often jumps between both oxygen atoms and the free energy barrier for proton exchange is very small (< 1 kcal/mol) (Supplementary Figure 9), consistent with the results of the reaction simulation. Therefore, the reaction products feature a low energy barrier hydrogen bond between Y223 and the new reducing end. Most importantly, this agrees with the reported neutron diffraction structure of the reaction products, which shows partial protonation of Y220 in *Ps*Alg7C (Figure 5B).

## Discussion

The reaction mechanism of PLs has been described as a β-elimination reaction of the O’-C4 glycosidic bond, leading to the formation of the 4,5-unsaturated sugar at the non-reducing end of the released product^11,16^ (Figure 1B). However, no high-resolution structure with well-defined substrate positions has been reported, hampering an atomistic description of the molecular mechanism. Here we reported high resolution structures of enzyme-substrate complexes of two PL7 that degrade alginates, *Ps*Alg7A and *Ps*Alg7C from the marine fungus *P. salina*. We showed by time-resolved NMR that the enzymes are endolytic, M specific and produce mainly trimers or oligomers with unsaturated ends. The structures include the first high resolution Michaelis-Menten complex of a PL7 captured using a single mutant (Y to F), minimally perturbing the active site architecture and, most importantly, with the substrate in a catalytically competent position. Together with a neutron diffraction structure, we uncovered the protonation states of the putative catalytic residues Y223/Y220 and H124/H229 (*Ps*Alg7A/*Ps*Alg7C numbering). We showed that both residues are unprotonated, suggesting that only Y223/Y220 is catalytically relevant. The crystal structures further revealed an active site groove with a complex hydrogen bond pattern likely optimized for alginate recognition. In addition, the first SBS observed on a PL was found on *Ps*Alg7C.

The crystal structure of the Michaelis-Menten complex of *Ps*Alg7A in complex with HexaM was used for QM/MM simulations of the reaction mechanism, which confirmed the β-elimination reaction and solved the complete mechanism at atomic detail. The reaction takes place in one-step (*i.e.,* a concerted process), but it is highly asynchronous, with proton abstraction by Y223 taking place before the TS. This is supported by the fact that Y220 is deprotonated in the neutron structure of *Ps*Alg7C but it is partially protonated after catalysis (the exchanged deuterium occupancy was 0.73). Our findings of a concerted yet asynchronous mechanism reconciles previous experimental studies which were interpreted in terms of either concerted or step-wise reactions. Our simulations also reveal the carbanion nature of the TS and the occurrence of a low energy barrier hydrogen bond between Y223 and the new reducing end at the reaction products.

The *+1* sugar adopts a relaxed chair conformation (^4^*C*_1_) at the Michaelis-Menten complex, but evolves gradually towards ^2^*S*_O_ at the TS, thanks to a subtle but important motion of Y223 and R82, delineating a ^4^*C*_1_ → [^2^*S*_O_]^‡^ → ^2^*E* conformational itinerary. Importantly, the conformations attained by the *+1* sugar during the reaction lie in a low energy region of the conformational FEL of the substrate at the Michaelis-Menten complex. This indicates that, similar to what has been observed in glycosidases,^1,2^ the active site of *Ps*Alg7A is preorganized for catalysis not only in terms of having the catalytic residues at the right position, but also by displaying a binding cavity in which *only* catalytic conformations fit optimally.

The finding of a concerted (one-step) reaction for *Ps*Alg7A clarifies previous findings of kinetic analyses for several PL7 enzymes, where the presence of a reaction intermediate remained ambiguous^16,40^. In the case of PL8 Chondroitin AC lyase from *Flavobacterium heparinum*, which also undergoes *syn*-β-elimination^41^, previous KIE and LFER experiments suggested a step-wise reaction without glycosidic bond breaking in the rate-determining step. However, a semiempirical QM/MM study on a family PL8 hyaluronate lyase ^42^ found a concerted *syn*-β-elimination reaction. We argue that the kinetic data can also be interpreted in terms of a concerted reaction in which proton abstraction, and not glycosidic bond cleavage, occurs first, as we have found for PL7 (the reaction is highly asynchronous). Structural and kinetic experiments on other PL families such as PL1 and PL10, which undergo anti-elimination, support catalysis through either a stepwise E1cb reaction or a concerted asynchronous E2 reaction^43^, consistent with our observations. In both cases, a carbanion species was predicted to be formed, consistent with our results for *Ps*Alg7. Integrating our novel findings with existing literature, we propose that the concerted but asynchronous reaction that we have found for PL7 (Figure 7B) may extend to other PL families. These structural and mechanistic insight presented mark a significant advance in our understanding of alginate lyase mechanisms, providing avenues for the engineering of ALs for the production of tailor-made alginate oligosaccharides. Additionally, they open up opportunities for designing activity-based probes to identify alginate lyases in fungal and bacterial secretomes.

## Data Availability Statements

The X-Ray structures of PsAlg7A, PsAlg7A-hexaM, PsAlg7A-pentaM, PsAlg7A-tetraM, PsAlg7A-triM, PsAlg7A-diM, PsAlg7A-Y223F, PsAlg7A-Y223F-hexaM, PsAlg7A-Y223F-pentaM, PsAlg7A-Y223F-tetraM, PsAlg7A-Y223F-triM, PsAlg7A-Y223F-diM, PsAlg7C, PsAlg7C-hexaM, PsAlg7C-pentaM, PsAlg7C-tetraM, PsAlg7C-triM, PsAlg7C-diM, PsAlg7C-Y220F, PsAlg7C-Y220F-hexaM, PsAlg7C-Y220F-pentaM, PsAlg7C-Y220F-diM, PsAlg7C-H110N, PsAlg7C-H110N-hexaM, PsAlg7C-H110N-pentaM, PsAlg7C-H110N-tertaM and PsAlg7C-H110N-triM were deposited in the RCSB PDB with accession codes 6YWF, 7P25, 7ORY, 7P90, 7OOF, 7PBF, 7NL3, 7NCZ, 7NPP, 7NY3, 7O6H, 7NM6, 8C3X, 8PC8, 8PC3, 8PCX, 8PED, 8PDT, 8C0M, 8BJO, 8BXZ, 8P6O, 8RBI, 8QMJ, 8QIZ and 8QLI, 8R43, respectively. Joint X-ray and neutron diffraction structures of PsAlg7C and PsAlg7C-pentaM were deposited in the RCSB PDB with accession codes 8RBR and 8RBN, respectively. Data files of the classical MD simulation and QM/MM OPES simulations have been deposited in Zenodo and will be released upon publication.

## Acknowledgements

This study was funded by the European Commission JPI Cofund Blue BioEconomy *via* the MARIKAT project (9082-00021B), the Research Council of Norway (RCN) *via* the AlgModE project (315385), the Independent Research Fund Denmark *via* the DSMADE project (DFF170746) and by the Department of Biotechnology and Biomedicine (DTU Bioengineering), Enzyme Discovery & Engineering Program, Technical University of Denmark, the Spanish Ministry of Science and Innovation (MICINN/AEI/FEDER, UE, PID2020-118893GB-100, to CR), the Spanish Structures of Excellence María de Maeztu (CEX2021-001202-M, to CR), the Agency for Management of University and Research Grants of Catalonia (AGAUR, 2021-SGR-00680, to CR) and the European Research Council (ERC-2020-SyG-951231 “Carbocentre”, to CR). JPR was supported by MICINN (FPI fellowship PRE2021-097898). FF and DHW were supported by The Novo Nordisk Foundation through grant number NNF10CC1016517 to the Novo Nordisk Foundation Center for Biosustainability at the Technical University of Denmark, which also supported the collection of some of the crystallographic data. We thank the Danish Agency for Science, Technology, and Innovation for funding the instrument center DanScatt, which supported our usage of beam time at the synchrotrons. We acknowledge MAX IV Laboratory for time on Beamline BioMAX under Proposal MX20190334 and MX20200120. Research conducted at MAX IV, a Swedish national user facility, is supported by the Swedish Research council under contract 2018 - 07152, the Swedish Governmental Agency for Innovation Systems under contract 2018-04969, and Formas under contract 2019-02496. We acknowledge synchrotron beamline P13 and P14 operated by EMBL Hamburg at the PETRA III storage ring (DESY, Hamburg, Germany), on which data were also collected. We acknowledge the European Synchrotron Radiation Facility for provision of beam time on ID30B operated by EMBL Grenoble. We acknowledge the Spallation Neutron Source, a DOE Office of Science User Facility operated by the Oak Ridge National Laboratory for provision of beam time. RCN is also acknowledged for funding the Norwegian NMR Platform (226244), SBP-N (294946) and the Carlsberg Foundation for funding the Mosquito crystallization robot, which was used for obtaining some of the protein crystals. We are grateful for the technical assistance provided by the support of the MareNostrum IV and CTE-Power supercomputers of the Barcelona Supercomputing Center (BSC-CNS), within the Red Española de Supercomputación (RES).

## Author Contribution

C.W. and C.R. conceived and designed the study with input from A.S.M.; M.V., B.P., L.J.K, C.L.-H. performed all molecular biology and biochemical experiments with help from F.L.A. and C.W.; C. W. and F. F. performed all the X-ray crystallography experiments with help from D. H. W.; C.W., J.P.M. and F.M. performed the neutron crystallography experiments; J.P.R.-F. performed the computational experiments. C.W. and C.R. wrote the article, with contributions from all authors. All authors read and approved the final manuscript.

## Online methods

### Protein production and site-directed mutagenesis

*Ps*Alg7A-Y223F, *Ps*Alg7C-H124N and *Ps*Alg7C-Y220F were constructed by site-directed mutagenesis using CloneAmp polymerase (Takara), a set of mutagenic primers (see Table S4), and pPICzαA/PsAlg7A or pPICzαA/PsAlg7C as template. Plasmid templates were then digested with DpnI at 37°C overnight, and the resulting PCR products were purified using the illustra GFX PCR DNA and Gel Band Purification Kit (GE Healthcare Life Sciences). The purified PCR products were subsequently transformed into *Escherichia coli* DH5α and plated on LB low salt agar supplemented with Zeocin (25 µg ml^-1^). Positive transformants were selected, and the corresponding plasmids were extracted using the GeneJET Plasmid Miniprep Kit (ThermoFisher Scientific). The validity of all constructs was confirmed through sequencing (Macrogen). Cloning into *Pichia pastoris* X-33 was carried out following the established protocol as in Pilgaard *et al.*^1^.

Expression and purification of *Ps*Alg7A (GenBank acc. nr. VFY81779.1), *Ps*Alg7A-Y223F, *Ps*Alg7C (GenBank acc. nr. CAD6594633), *Ps*Alg7C-H124N and *Ps*Alg7C-Y220F where carried out as in Pilgaard *et al.*^1^ except that a size exclusion chromatography step on a HiLoad 16/60 Superdex G75 (GE Healthcare) equilibrated with 10 mM Tris-HCl, 150 mM NaCl pH 7 followed the previous purification steps with a flow of 0.5 ml min^-1^. The purity of the obtained fractions were evaluated on 12% SDS-PAGE gels and the pure fractions were pooled and stored at 4 °C.

### Time resolved NMR

Each reaction was comprised of 160 µL of 10 mg/mL ^13^C-1-enriched polymannuronate substrate^2^ in buffer (with 99.9% D_2_O) and 10 µL of enzyme stock. For *Ps*PL7A (enzyme stock 147.5 µg/mL) the buffer composition was 5 mM NaOAc pD 5.0, 10 mM NaCl, 1.5 mM ZnCl_2_ and for *Ps*PL7C (enzyme stock 150 µg/mL) the buffer composition was 5 mM HEPES pD 8.0,

100 mM NaCl. All experiments were recorded on Bruker 800 MHz Avance III HD spectrometer (Bruker BioSpin AG, Fällanden, Switzerland) equipped with a 5 mm cryogenic CP-TCI z-gradient probe at 25°C.

A ^13^C-monitored psuedo-2D spectrum followed by a ^13^C-^1^H HSQC (heteronuclear single quantum coherence) spectrum was recorded for each reaction. The psuedo-2D consists of a series of 1D carbon spectra recorded every 5 minutes for a total of 200 spectra (total experiment time of 16h 40min). Each carbon spectrum (using inverse gated proton decoupling) contains 32 K data points, spectral width of 200 ppm, 32 scans with a 30° flip angle and relaxation delay of 2.1 s (total recording time of 81 s per spectrum). Signals were assigned based previously published assigments from Heyraud *el al*., (1996)^3^, Li *et al.* (2011)^4^, and Mathieu *el al.* (2016)^5^. The spectra were recorded, processed, and analyzed using TopSpin 3.6pl7 and TopSpin 4.0.7 software (Bruker BioSpin AG, Fällanden, Switzerland).

### Crystallization, data collection and data processing for X-ray structures

*Ps*Alg7A (14 mg ml^-1^), *Ps*Alg7A-Y223F (15 mg ml^-1^), *Ps*Alg7C (7.5 mg ml^-1^), *Ps*Alg7C-H124N (7.5 mg ml^-1^) and *Ps*Alg7C-Y220F (7.5 mg ml^-1^) in 10 mM Tris-HCl, 150 mM NaCl pH 7 were crystallized in 48-well MRC Maxi plates (Jena Bioscience) (see crystallization conditions in Supplementary Table 2). The crystals soaked with di-, tri-, tetra-, penta- or hexa-mannuronic sodium salt (Carbosynth) by adding a few crystals to the drop that were then allowed to equilibrate for 1–24h. The crystals were supplemented with PEG1500 for cryoprotection, and the crystals were cryocooled in liquid nitrogen. Data were collected at BioMAX^6^ at the MAX IV Laboratory and ID30B^7^ at the European Synchrotron Radiation Facility using MxCUBE3^8^ and P13^9^ and P14 at PETRA III using MxCUBE2^10^. The dataset were processed to 0.82–2.00 Å resolution, respectively using XDSapp^11^, Xia2^12^ autoPROC^13^ or EDNA^14^ (see Supplementary Table 2 for statistics) in space groups *P*1, *P*12_1_1, *P*2_1_2_1_2_1_ or *C*12_1_.

### Phasing and refinement of X-ray structures

The structure of *Ps*Alg7A was determined by molecular replacement with PHASER^15^ from the Phenix package^16^ using the closest homolog at that time from the Protein Data Bank (www.pdb.org)^17^ as template, which was the PL7 alginate lyase A1-II’ from *Sphingomonas* sp. A1 (PDB ID 2CWS)^18^ identified through PDB-BLAST^19^ with a coverage around 65% and identity of around 28%. The initial structure was built with PHENIX.autobuild^20^ and then refined with PHENIX.refine^21^ and manual model rebuilding in Coot^22^. The other *Ps*Alg7A and *Ps*Alg7C structures were obtained similarly but using *Ps*Alg7A as the template. The sugar moieties conformations were analyzed with Privateer^23^.

### Crystallization, data collection at room temperature and data processing for neutron structures

*Ps*Alg7C (15 mg ml^-1^) in 10 mM Tris-HCl, 150 mM NaCl pH 7 was crystallized in Linbro plates (Jens Bioscience) by mixing 12 µl protein with 6 µl 0.1 M Bis-Tris pH 5.5, 30% PEG3350 and 0.3 M NaCl dissolved in D_2_O and incubated with 200 µl 0.1 M Bis-Tris pH 5.5, 30% PEG3350 and 0.3 M NaCl dissolved in H_2_O in the reservoir. After one-week crystals appeared and the buffer was then exchanged by removing 5 µl followed by addition of 5 µl 0.1 M Bis-Tris pH 5.5, 30% PEG3350 and 0.3 M NaCl dissolved in D_2_O and incubation for 24 h. This was repeated 5 times. Hexa-mannuronic sodium salt (Carbosynth) was added by dropping a few crystals on the drop that were then allowed to equilibrate for one week. The crystals were mounted in thin-walled quartz capillaries (Hampton) and sealed with beeswax.

Neutron time-of-flight diffraction data were collected at room temperature on the MaNDi instrument at the SNS^24,25^. An incident neutron wavelength bandpass of 2–4 Å was used. For the *Ps*Alg7C, a total of 7 diffraction images were collected with a Δφ of 10° and an average exposure of 24h per frame. Following neutron diffraction data collection, an X-ray dataset was collected on the same crystal at room temperature on a microfocus rotating anode X-ray diffractometer. A total of 360 diffraction images were collected with a Δφ of 1.0° and an exposure of 7 s per frame.

For *Ps*Alg7C in complex with PentaM, a total of 11 diffraction images were collected with a Δφ of 10° and an average exposure of 36 h per frame. Following neutron diffraction data collection, an X-ray dataset was collected on the same crystal at room temperature on a microfocus rotating anode X-ray diffractometer. A total of 180 diffraction images were collected with a Δφ of 1.0° and an exposure of 30 s per frame.

The neutron datasets were reduced using the *Mantid* package^26^ and integrated using three-dimensional profile fitting^27^. The data were wavelength normalized using *LAUENORM* from the *LAUEGEN* suite^28–30^. The X-ray data were indexed and integrated using *CrysAlis PRO* (Rigaku, Woodlands, Texas, USA) and scaled and merged with *AIMLESS* in the *CCP4* suite^31^. X-ray and neutron data collection statistics are presented in Supplementary Table 2.

### Phasing and refinement of neutron structures

The structures of *Ps*Alg7C were determined by molecular replacement with PHASER^15^ from the Phenix package^16^ using *Ps*Alg7C (PDB ID 8C3X) as the template. Joint X-ray/neutron refinement was performed using the Phenix software suite^16^ and manual model rebuilding in Coot^22^ (see Supplementary Table 3 for statistics).

### Molecular dynamics simulations

The starting structure for all molecular dynamics (MD) simulations conducted in this study was the crystal structure of *Ps*Alg7A-Y223F in complex with HexaM (PDB ID 7NCZ). The catalytic residue mutation (Y223F) was reverted using PyMOL (Schödinger). The simulations were performed under conditions of pH 5 and 200 mM NaCl, which represent the optimal conditions for in vitro enzymatic analysis^32^. All arginine and lysine residues were positively charged, while all aspartate and glutamate residues, as well as all carboxylate groups of the M moieties, were negatively charged. The protonation states of the histidine residues were chosen using the H++ server^33^ and guided by experimental data. The hydroxyl group of the Y223 catalytic residue was considered to be deprotonated, as per the β-elimination mechanism proposed in the literature and in previous works on related enzymes^34,35^. The system was positioned at the centre of a periodically repeated rectangular box (77.725 x 83.850 x 90.199 Å^3^), with a minimum distance of 15 Å between the solute surface and the box edge. The box was solvated with 14859 TIP3P^36^ water molecules and sodium and chlorine ions were added to mimic the salt concentration for optimal activity (200 mM). The protein was described using the Amber ff14SB force field^37^, while the GLYCAM06 force filed^38^ was used to describe the carbohydrate molecule. Atomic partial charges (RESP) of Y223 (negatively charged) were calculated at the HF/6-31G* level of theory using Gaussian09^39^. The topology and coordinate files for the classical MD simulations were generated using the LEaP module of AmberTools21^40^. The simulations were performed using Amber20 ^40^ with the CUDA version of the PMEMD module^41^. The solvated system was subjected to an energy minimization protocol, wherein the solvent was minimized first with 5000 steps of steepest descent minimization and 5000 steps of conjugate gradient minimization, with positional restraints applied to all atoms of the protein and the ligand. Subsequently, the entire system was minimized with 15000 steps of steepest descend and conjugate gradients (7500 each). The system was then gradually heated to 308 K in the NVT ensemble using the Berendsen thermostat, with a time constant of 1.0 ps for the coupling and 50 kcal mol^-1^Å^-2^ positional restraints applied to the heavy atoms of the protein and the substrate. The restraints were gradually decreased to 5 kcal mol^-1^Å^-2^ over three 500 ps steps, followed by 1000 ps of NPT equilibration using the Berendsen thermostat and barostat to maintain the system at 308 K and 1 atm, ensuring density equilibration. Three replicas were run for 250 ns, taking 50 ns as equilibration and 200ns as production (Supplementary Figure 6). The SHAKE algorithm^42^ was used to constrain all bonds involving hydrogen atoms and the time step was set to 2 fs.

Production runs in the NPT ensemble were controlled by the Langevin thermostat and the Berendsen barostat.

### QM/MM MD simulations of the β-elimination reaction

QM/MM MD simulations were employed to investigate the reaction mechanism in a less computationally demanding manner. This method involves partitioning the system into Quantum Mechanics (QM) and Molecular Mechanics (MM) fragments, where the QM region were treated using Born-Oppenheimer Density Functional Theory (DFT) based MD simulation, while the MM region were treated using force-field strategy. A snapshot form the classical MD trajectory (Supplementary Figure 6) was used as the starting point for the QM/MM MD simulations. A large QM region (112 atoms) was selected, including the sidechains of residues R82, Q127, K219 and Y223, the saccharide residues at the *+1* and *-1* subsites and part of the ones at the *+2* and *-2* subsites (Supplementary Figure 8). The QM region was enclosed in a cell of size 17.63 × 17.68 × 23.39 Å^3^, and the atoms at the QM -MM border were saturated with hydrogen atoms using the IMMOM approach^43^. CP2K v9.0^44,45^ was used for the QM/MM MD simulations. The QM region was treated at the DFT (PBE) level, employing the dual basis set of Gaussian and plane-waves (GPW) formalism. The wave function was expanded using the Gaussian triple- ζ valence polarized (TZV2P) basis set, and the electron density was converged using the auxiliary plane wave basis set with a density cut-off of 330 Ry and GTH pseudopotentials^46^. After relaxation, the system was equilibrated for 7 ps. Subsequently, the On-the-fly Probability Enhanced Sampling (OPES) method^47^ in its exploratory variant (OPES_E_) was used to explore the Free Energy surface. All QM/MM MD simulations were performed in the NVT ensemble using a coupling constant of 10 fs and an integration time step of 0.5 fs. Two CVs were used to activate the chemical reaction (Figure 6): the first CV (CV1) was taken as the distance of the scissile glycosidic bond (O4’-C4). The second CV (CV2) was taken as a linear combination of distances involving H5 during the reaction, with negative values indicating the absence of a bond and positive values indicating its presence in the reactant state. The time interval between two consecutive kernels was set to 50 fs. To ensure a better description of the transition state (TS) and prevent exploration of non-catalytic high-energy regions, we set the initial OPES_E_ energy barrier to 30 kcal mol^-1^. The simulation was stopped after two recrossing over the TS (Supplementary Figure 10). The minimum energy path was determined using the MEPS software^48^, and the TS was further refined using iso-committor analysis (Supplementary Figure 11). The MD trajectories generated in the simulations were analyzed using VMD^49^, and distances between atoms were calculated using the driver module of PLUMED^50^. Plots were generated using Matplotlib^51^, and figures of molecular structures were rendered using PyMOL (Shrödinger).

### QM/MM conformational analysis of sugar +1

To mitigate any potential bias inherent in the classical runs, we employed QM/MM metadynamics simulation to investigate the conformational landscape of the subsite +1 sugar of the ManAC_7_ substrate within the enzyme (Supplementary Figure 7). The simulation was initiated using the previously mentioned relaxed QM/MM structure (refer to the QM parameters provided above). Collective variables based on Cremer-Pople puckering coordinates, specifically q_x_, q_y,_ and q_z_ divide by the amplitude Q, were utilized. Gaussian functions were employed for deposition, with a deposition pace of 60 MD steps (30 fs), and width of 0.05 Å, 0.05 Å and 0.03 Å for q_x_/Q, q_y_/Q and q_z_/Q, respectively. The height of the Gaussians varied from 0.4 kcal/mol at the beginning of the simulation to 0.1 kcal/mol at the end. A total of 6789 Gaussians were deposited, and the simulation was carried out for a final duration of 203.65 ps.

Coordinate files and other simulation data can be found in the Zenodo repository (https://doi.org/10.5281/zenodo.10213646).

### Structural analysis

Structural alignments were obtained using PyMOL v. 2.5.4 (Schrödinger, LLC, New York; also used for rendering structural models). Electrostatic maps were obtained with the APBS-PDB2PQR software suite (https://server.poissonboltzmann.org/) using APBS v. 3.4.1 and PDB2PQR v. 3.6.1^52^. Hydrophobicity plots were obtained using the color_h.py script in PyMOL v. 2.5.4 (Schrödinger) and colored according to the Eisenberg hydrophobicity scale^53^. ProBiS H_2_O was used to identify conserved water molecules^54^. LigPlot+ v.2.2^55^ and PyMOL v. 2.5.4 (Schrödinger) was used to analyze hydrogen bonds.

