## Supplementary Information for "Unraveling the Molecular mechanism of Polysaccharide Lyases for Efficient Alginate Degradation"

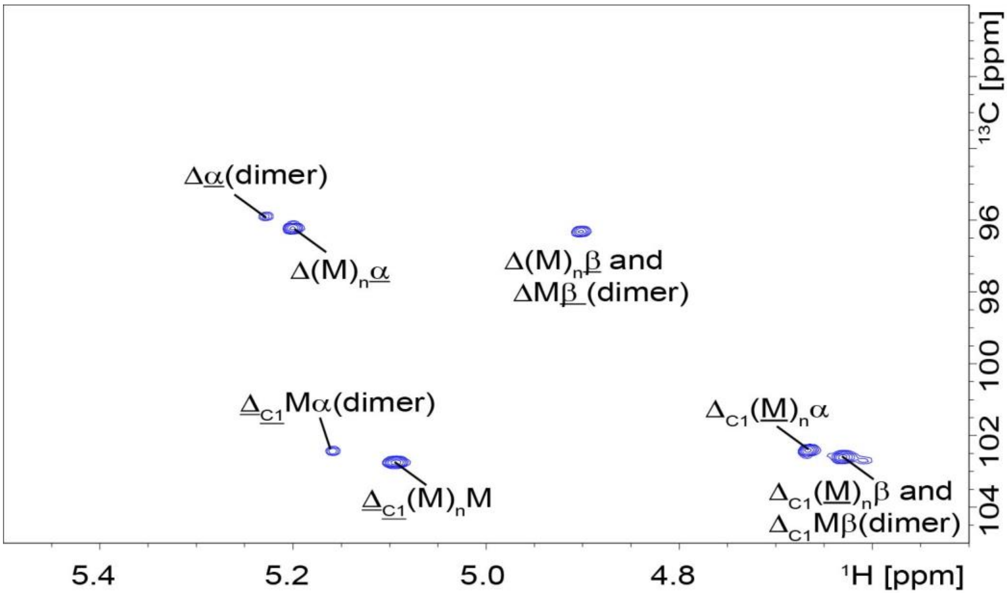


**Supplementary Figure 1**. **$^{1}$H-$^{13}$C HSQC spectrum anomeric region after treatment of $^{13}$C-1-enriched-polyM treated with *Ps*PL7A**. Reaction was completed within a 3mm NMR tube with 160 µL of substrate buffer solution (10 mg/mL $^{13}$C-1-polyM substrate in 5 mM HEPES, pD 8.0 with 100 mM NaCl in 99.9% $D_2O$) and 10 µL of 147.5 µg/mL *Ps*PL7A incubated at 25 °C. α and β indicate the signals for the reducing end of alginate, M is mannuronate, C1 indicate anomeric in alginate sugar ring, Δ: 4,5-unsaturated 4-deoxy-L-erythro-hex-4-enepyranosyluronate, underlined indicate the residue with the anomeric C1 giving rise to the signal.

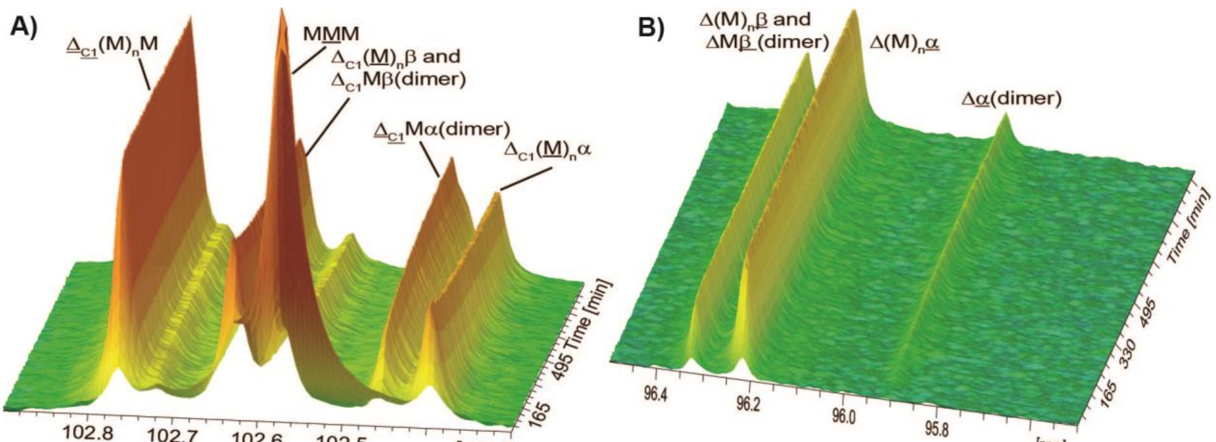


**Supplementary Figure 2: Time-resolved $^{13}$C-NMR spectra of $^{13}$C-1-enriched polyM treated with *Ps*PL7C.** Reaction was conducted in 3mm NMR tube with 160 µL of substrate buffer solution (10 mg/mL $^{13}$C-1-polyM substrate in 5 mM Na-acetate, pD 5.0 with 10 mM NaCl and 1.5 mM $ZnCl_2$ in 99.9% $D_2O$) and 10 µl of 150 µg/mL *Ps*PL7C incubated at 25 °C. Pseudo 2D time-resolved spectra recorded by acquiring 1D carbon spectra every 5 mins for a total of 16 hrs 40 mins. Panel A) shows anomeric region and B) reducing end region in the carbon spectra. α and β indicate the signals for the reducing end of alginate, M is mannuronate, C1 indicate anomeric in alginate sugar ring, Δ: 4,5-unsaturated 4-deoxy-L-*erythro*-hex-4-enepyranosyluronate, underlined indicate the residue with the anomeric C1 giving rise to the signal.

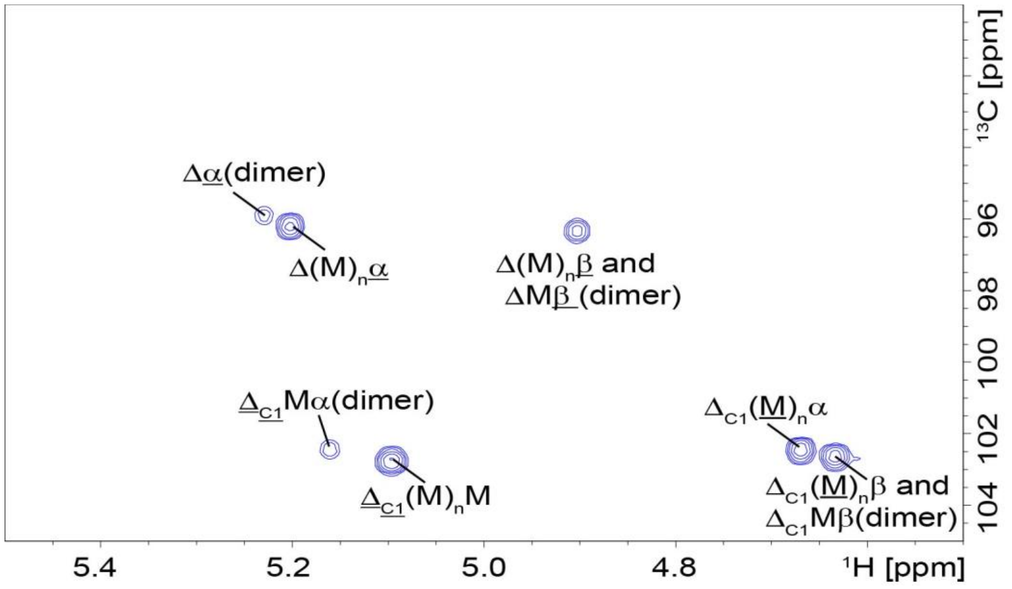


**Supplementary Figure 3**. **$^{1}H$-$^{13}C$ HSQC spectrum anomeric region after treatment of $^{13}C$-1-enriched-polyM treated with *Ps*PL7C**. Reaction was completed within a 3mm NMR tube with 160 μL of substrate buffer solution (10 mg/mL $^{13}C$-1-polyM substrate in 5 mM Na-acetate, pD 5.0 with 100 mM NaCl and 1.5 mM $ZnCl_2$ in 99.9% $D_2O$) and 10 μL of 150 μg/mL *Ps*PL7C incubated at 25 °C. α and β indicate the signals for the reducing end of alginate, M is mannuronate, C1 indicate anomeric in alginate sugar ring, Δ: 4,5-unsaturated 4-deoxy-L-erythro-hex-4-enepyranosyluronate, underlined indicate the residue with the anomeric C1 giving rise to the signal.

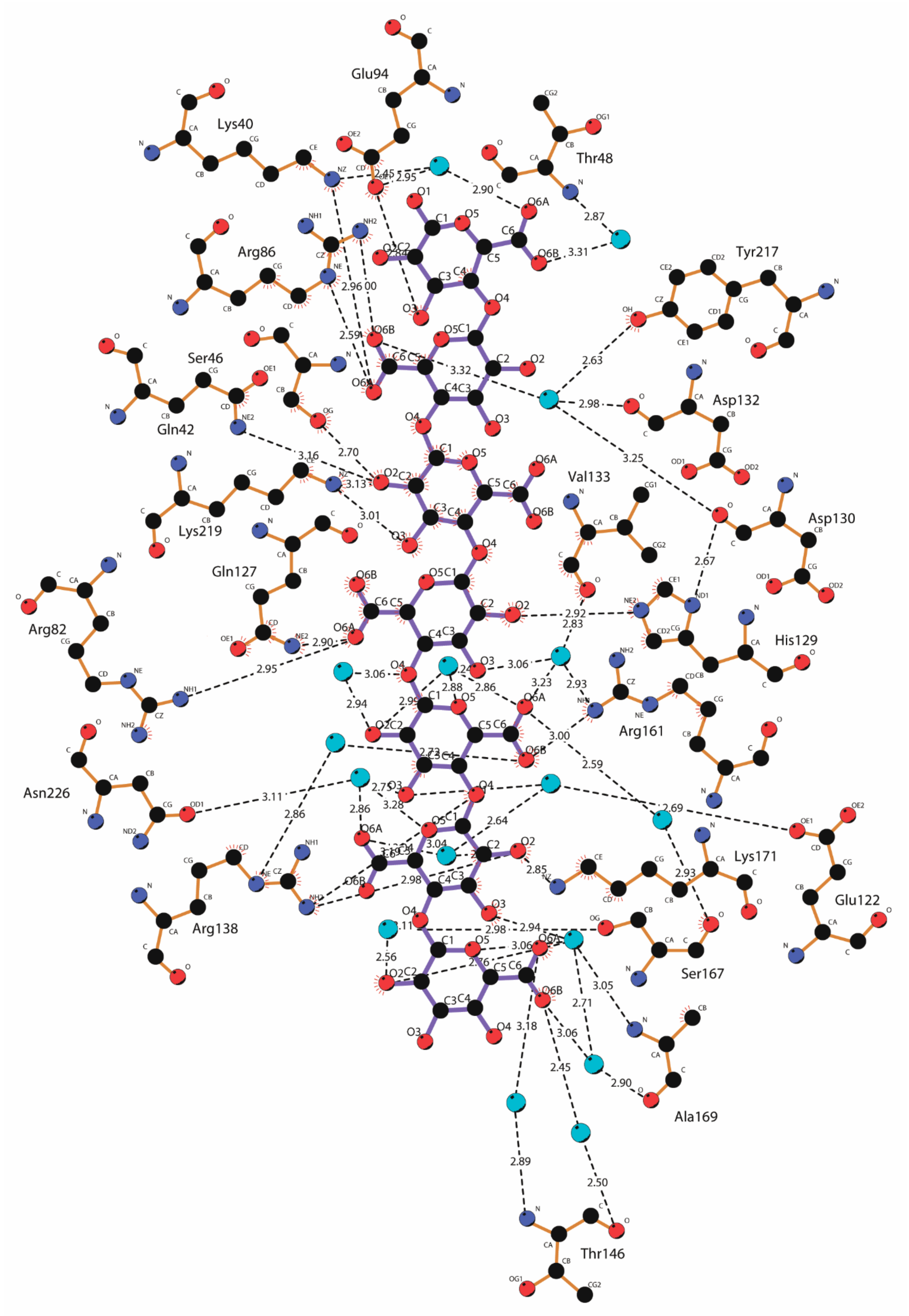


**Supplementary Figure 4.** Putative hydrogen bonds between β-D-mannuronic acid moieties and *Ps*Alg7A-Y223F visualized with LigPlot+[55].

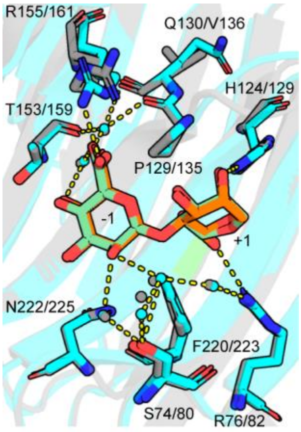


**Supplementary Figure 5.** Conserved water molecules and residues and the putative hydrogen bonds formed to the M moieties at subsites *-1* and *+1* for PsAlg7A-Y220F (grey) in complex with HexaM (green) and PsAlg7C (cyan) in complex with HexaM (orange). Hydrogen bonds are shown as dotted lines (yellow).

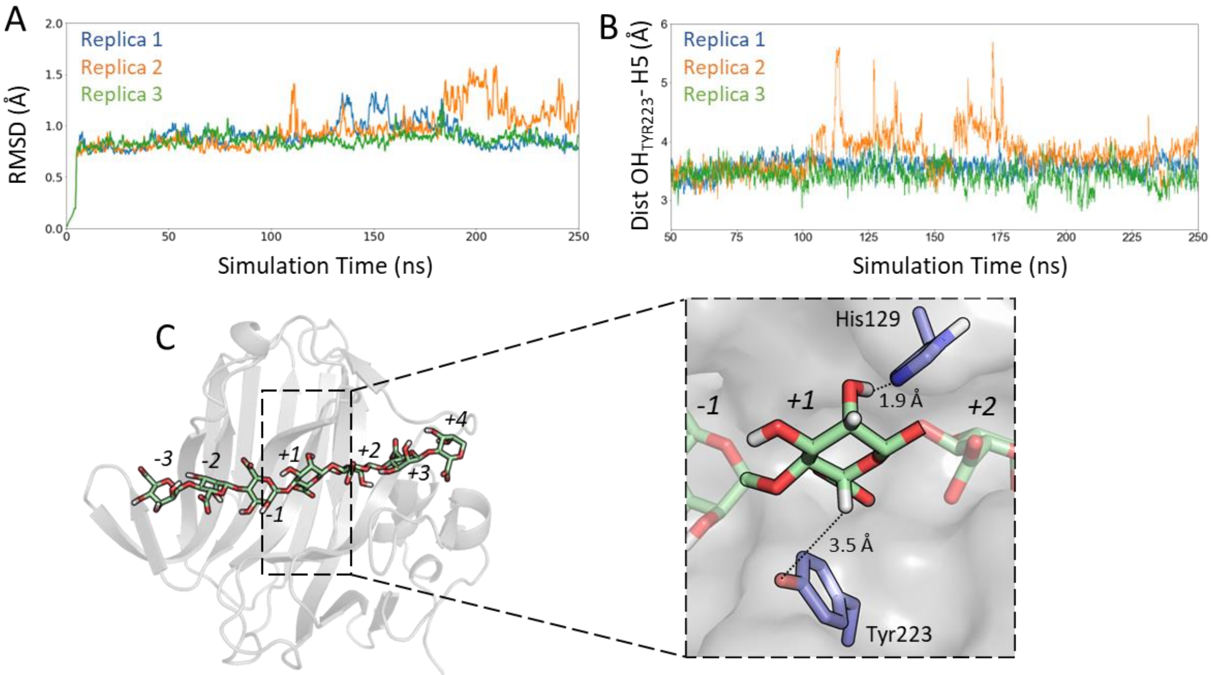


**Supplementary Figure 6. Classical MD simulations of PsPL7A in complex with heptamannuronic acid**. A) Time evolution of the protein backbone RMSD. The 250ns simulated for each of the replicas are represented independently using blue, orange and green colors for the replica 1, 2 and 3 respectively. B) Time evolution of the catalytic distance $OH_{Tyr223}$ – H5. The 200ns of production is presented for each of the replicas using the same colours as before. C) Michaelis-Complex representations of the snapshot taken from the classical MD and used for the QM/MM simulations with zoom-in in the +1 subsite showing how sugar +1 is sandwiched by residues Tyr223 and His129.

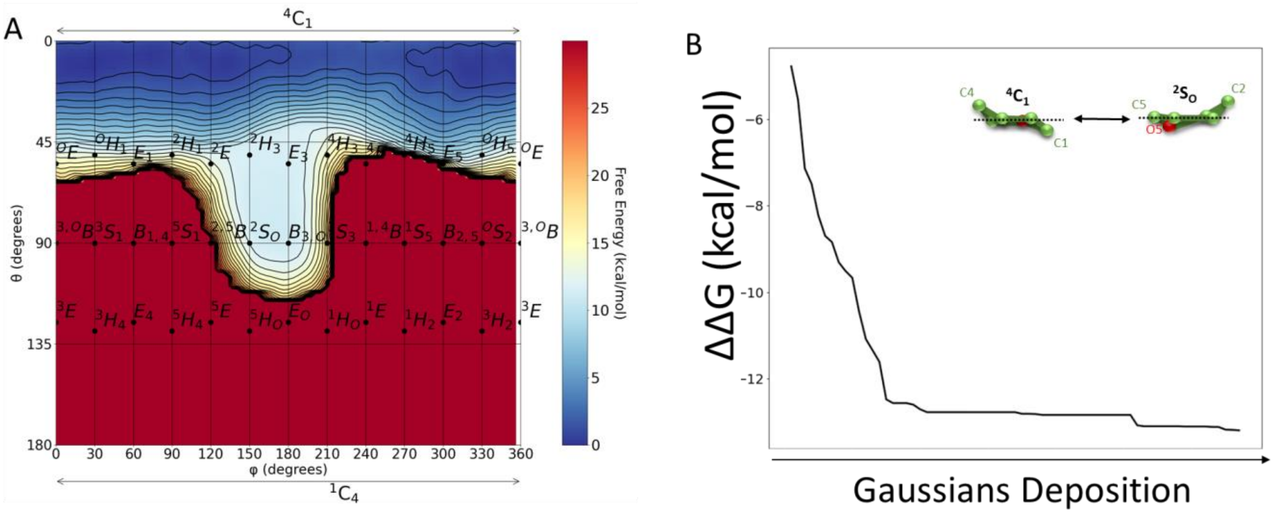


**Supplementary Figure 7. Conformational landscape of β-mannurosyl unit *+1*.** A) Cremer-Pople Mercator representation of the FEL obtained using QM/MM Metadynamics. The global minimum corresponds to the $^{4}C_{1}$ conformation, with a low-energy region that extends towards $^{2}S_{O}$. B) Evolution of the barrier between $^{4}C_{1}$ and $^{2}S_{O}$ conformation with new gaussian depositions to assess convergence in the QM/MM metadynamic simulation.

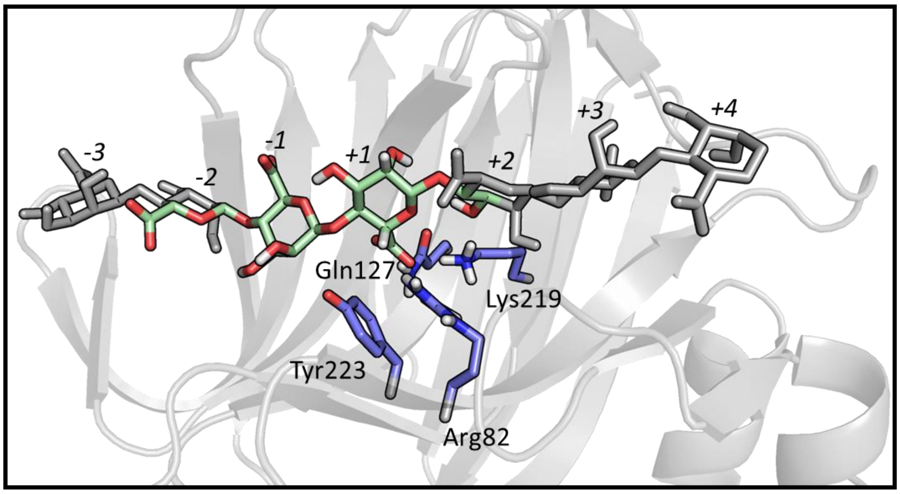


**Supplementary Figure 8. Scheme with stick representation of the atoms included in the QM region for modelling the reaction**. QM atoms are colored while MM atoms are presneted in dark grey. Labels for both, aminoacids and sugar units, are shown.

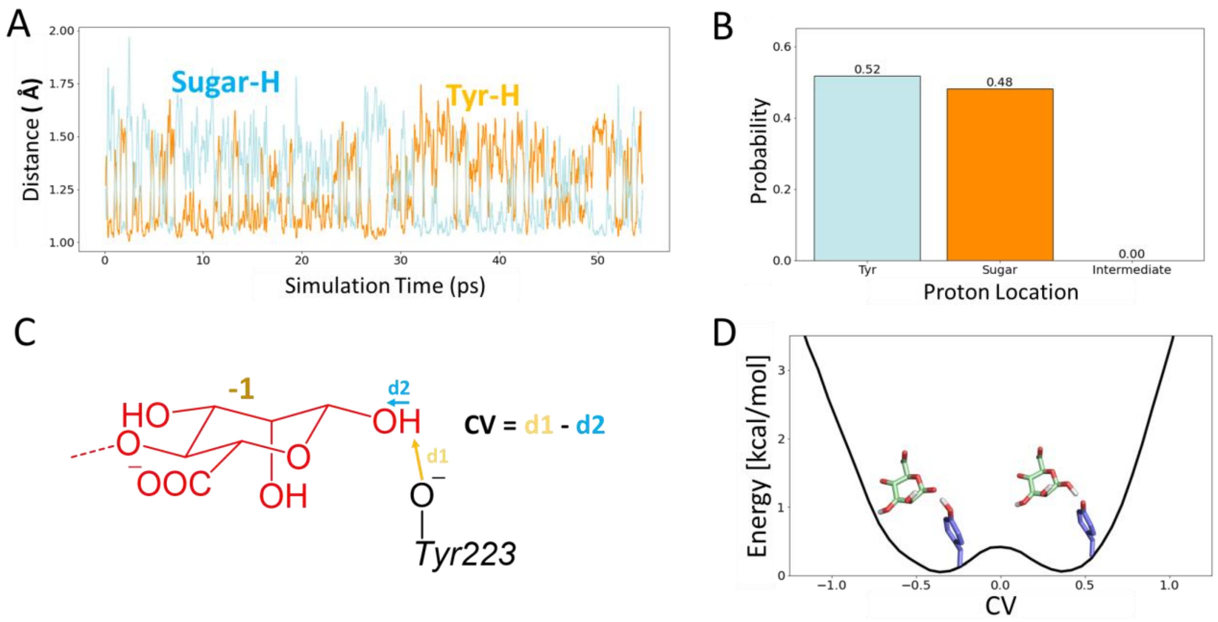


**Supplementary Figure 9. Proton transfer analysis at product state using QM/MM simulation**. A) Comparison between distances catalytic proton – new reduced end (blue) and catalytic proton – catalytic tyrosine (orange). B) Probability distribution of proton location. C) CV used for reconstructing the proton transfer FES. D) FES with proton location representation.

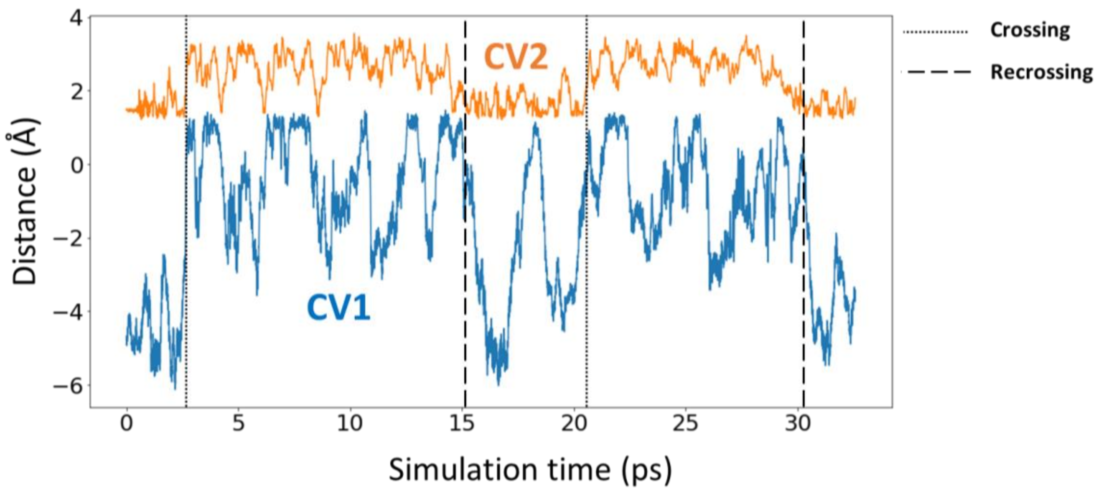


**Supplementary Figure 10. Time evolution of the CVs used for driving the reaction in the QM/MM OPES$_E$ simulation**. CV1 and CV2 are colored in blue and orange respectively. Dotted line is used to highlight a crossing, while a dash line donates a recrossing. The first recrossing is reach after 15 ps of simulation, while the second is observed after 30 ps. Both CVs are explained in the section QM/MM MD simulations of the β-elimination reaction and schematized in Fig. 7 in the main text.

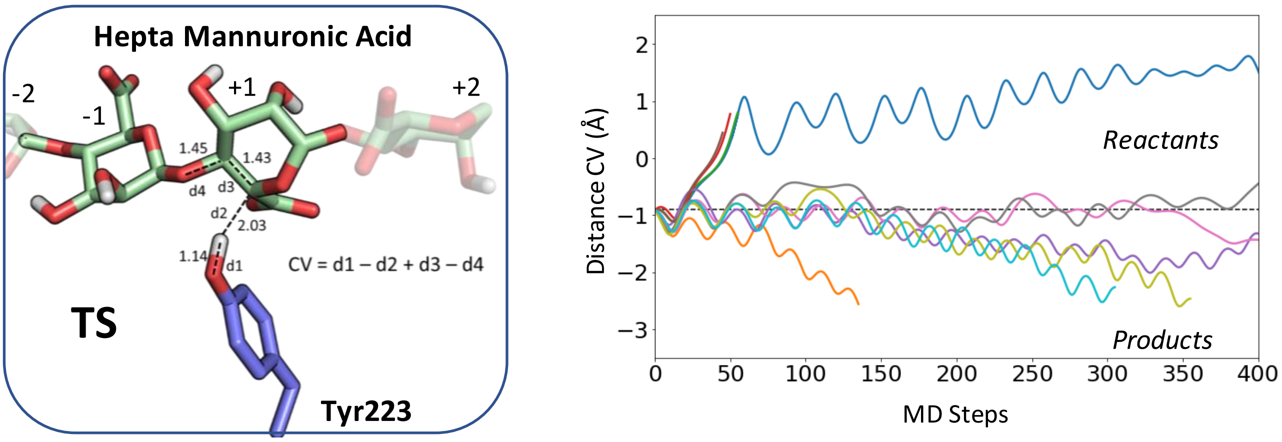


**Supplementary Figure 11. Iso-committor analysis**. Candidate TS structure used for the 10 QM/MM runs with different initial velocities is shown on the left, with distances in angstroms. The CV (linear combination of catalytically relevant distances) to follow the evolution is also described. On the right, time evolution of the CV for each of the 10 independents QM/MM MD runs. A dashed line highlights the value of the CV at the initial structure; values larger/smaller than 0.93 Å indicate simulations evolving towards reactants/products, respectively.

**Supplementary Table 1**: X-ray data collection and refinement statistics.

| Enzyme | ***Ps*Alg7A** | ***Ps*Alg7A** | ***Ps*Alg7A** | ***Ps*Alg7A** | ***Ps*Alg7A** |
|---|---|---|---|---|---|
| Ligand | - | **HexaM** | **PentaM** | **TetraM** | **TriM** |
| PDB ID | 6YWF | 7P25 | 7ORY | 7P90 | 7OOF |
| Crystallization condition | 20% PEG 1500 and 0.1 M TBG buffer pH 4 | 20% PEG 1500 and 0.1 M TBG buffer pH 4 | 20% PEG 1500 and 0.1 M TBG buffer pH 4 | 20% PEG 1500 and 0.1 M TBG buffer pH 4 | 20% PEG 1500 and 0.1 M TBG buffer pH 4 |
| Protein:reservoir ratio | 2:1 | 2:1 | 2:1 | 2:1 | 2:1 |
| Cryoprotectant | PEG1500 | PEG1500 | PEG1500 | PEG1500 | PEG1500 |
| Beamline | P14 | BioMax | BioMax | BioMax | BioMax |
| **Data collection** | | | | | |
| Wavelength (Å) | 0.98 | 0.98 | 0.95 | 0.95 | 0.95 |
| Resolution (Å) | 40.22–1.90 (1.97 –1.90) | 37.15–1.47 (1.50–1.47) | 40.44–1.46 (1.51–1.46) | 36.41–1.87 (1.94–1.87) | 36.24–1-30 (1.35–1.30) |
| Space group | $P2_12_12_1$ | $P2_12_12_1$ | $P2_12_12_1$ | $P2_12_12_1$ | $P2_12_12_1$ |
| Unit cell | | | | | |
| *a*, *b*, *c* (Å) | 37.8, 80.4, 81.4 | 34.6; 74.3; 79.9 | 34.8; 80.9; 81.2 | 35.2; 80.9; 81.5 | 34.8; 81.1; 80.7 |
| Total no. of reflections | 157888 (11623) | 465918 (30921) | 496541 (29779) | 250050 (17943) | 711945 (43761) |
| No. of unique reflections | 18624 (1794) | 35825 (2367) | 40650 (3998) | 19631 (1698) | 56656 (5354) |
| $R_{merge}$ | 0.08 (0.33) | 0.12 (2.83) | 0.05 (1.68) | 0.05 (0.89) | 0.09 (2.01) |
| $R_{meas}$ | 0.09 (0.37) | 0.13 (2.95) | 0.06 (1.80) | 0.06 (0.93) | 0.09 (2.15) |
| $R_{pim}$ | 0.03 (0.14) | 0.03 (0.81) | 0.02 (0.63) | 0.02 (0.28) | 0.03 (0.73) |
| $CC_{1/2}$ | 0.99 (0.95) | 1.00 (0.63) | 1.00 (0.53) | 1.00 (0.89) | 1.00 (0.46) |
| $I/\sigma(I)$ | 7.09 (1.20) | 13.13 (1.39) | 18.05 (1.03) | 22.45 (2.43) | 15.45 (1.07) |
| Completeness (%) | 98.32 (93.09) | 99.91 (99.87) | 99.88 (99.33) | 98.39 (86.92) | 99.51 (95.42) |
| Redundancy | 8.5 (6.5) | 13.0 (13.1) | 12.2 (7.4) | 12.7 (10.6) | 12.6 (8.4) |
| **Refinement** | | | | | |
| Reflections used in refinement | 18372 (1712) | 35825 (2367) | 40622 (3986) | 19608 (1694) | 56789 (5353) |
| Reflections used for $R_{free}$ | 1830 (166) | 1107 (73) | 2032 (199) | 981 (85) | 2096 (198) |
| $R_{work}$ | 0.15 (0.18) | 0.17 (0.42) | 0.18 (0.40) | 0.17 (0.27) | 0.16 (0.35) |
| $R_{free}$ | 0.18 (0.22) | 0.19 (0.41) | 0.20 (0.39) | 0.21 (0.31) | 0.17 (0.35) |
| CC (work) | 0.96 (0.94) | 0.97 (0.84) | 0.96 (0.73) | 0.97 (0.92) | 0.42 (0.08) |
| CC (free) | 0.96 (0.93) | 0.96 (0.72) | 0.98 (0.74) | 0.98 (0.91) | 0.41 (0.11) |
| No. of refined non-hydrogen atoms | | | | | |
| Protein | 1737 | 1827 | 1782 | 1764 | 1811 |
| Ligand(s) | - | 73 | 86 | 129 | 106 |
| Solvent | 237 | 126 | 191 | 134 | 250 |
| *B*-factors | | | | | |
| Protein | 20.87 | 24.32 | 33.22 | 45.57 | 19.16 |
| Lignad(s) | - | 55.15 | 56.57 | 63.67 | 29.98 |
| Solvent | 31.88 | 31.49 | 40.58 | 48.03 | 30.51 |
| Wilson *B*-factor | 17.69 | 18.01 | 24.8 | 35.21 | 16.87 |
| r.m.s.d. | | | | | |
| Bond lengths (Å) | 0.003 | 0.01 | 0.01 | 0.01 | 0.01 |
| Bond lengths (°) | 0.70 | 0.93 | 1.40 | 1.13 | 1.00 |
| Ramachandran plot | | | | | |
| Favored | 97.31 | 96.44 | 96.88 | 95.09 | 97.32 |
| Allowed | 2.69 | 3.56 | 3.12 | 4.91 | 2.68 |
| Outliers | 0.00 | 0.00 | 0.00 | 0.00 | 0.00 |
| Rotamer outliers | 0.51 | 0.49 | 0.50 | 1.01 | 0.00 |
| No. of TLS groups | 8 | - | - | 3 | - |

**Supplementary Table 1**: X-ray data collection and refinement statistics.

| Enzyme | ***Ps*Alg7A** | ***Ps*Alg7A-Y223F** | ***Ps*Alg7A-Y223F** | ***Ps*Alg7A-Y223F** | ***Ps*Alg7A-Y223F** |
|---|---|---|---|---|---|
| Ligand | **DiM** | - | **HexaM** | **PentaM** | **TetraM** |
| PDB ID | 7PBF | 7NL3 | 7NCZ | 7NPP | 7NY3 |
| Crystallization condition | 20% PEG 1500 and 0.1 M TBG buffer pH 4 | 20% PEG 1500 and 0.1 M TBG buffer pH 4 | 20% PEG 1500 and 0.1 M TBG buffer pH 4 | 20% PEG 1500 and 0.1 M TBG buffer pH 4 | 20% PEG 1500 and 0.1 M TBG buffer pH 4 |
| Protein:reservoir ratio | 2:1 | 2:1 | 2:1 | 2:1 | 2:1 |
| Cryoprotectant | PEG1500 | PEG1500 | PEG1500 | PEG1500 | PEG1500 |
| Beamline | BioMax | BioMax | BioMax | BioMax | BioMax |
| **Data collection** | | | | | |
| Wavelength (Å) | 0.98 | 0.98 | 0.98 | 0.98 | 0.98 |
| Resolution (Å) | 40.32-1.54 (1.60-1.54) | 40.31–1.70 (1.76–1.70) | 40.45–1.64 (1.70–1.64) | 36.31–1.45 (1.50–1.45) | 40.47–1.82 (1.89–1.82 |
| Space group | $P2_12_12_1$ | $P2_12_12_1$ | $P2_12_12_1$ | $P2_12_12_1$ | $P2_12_12_1$ |
| Unit cell | | | | | |
| *a, b, c* (Å) | 35.0; 80.6; 81.2 | 34.8; 80.6; 80.9 | 34.8, 80.9, 81.2 | 34.8, 80.7, 81.3 | 80.9, 34.8, 81.0 |
| Total no. of reflections | 443443 (40888) | 331065 (29899) | 373874 (31064) | 279808 (15618) | 209775 (15296) |
| No. of unique reflections | 34394 (3256) | 25527 (2411) | 28633 (2528) | 41381 (3965) | 19973 (2048) |
| $R_{merge}$ | 0.08 (2.72) | 0.13 (1.76) | 0.16 (2.04) | 0.08 (1.06) | 3.81 (4.56) |
| $R_{meas}$ | 0.08 (2.84) | 0.14 (1.83) | 0.16 (2.12) | 0.09 (1.22) | 4.72 (6.01) |
| $R_{pim}$ | 0.02 (0.79) | 0.04 (0.51) | 0.05 (0.60) | 0.03 (0.59) | 2.58 (3.75) |
| $CC_{1/2}$ | 1.00 (0.55) | 1.00 (0.50) | 1.00 (0.55) | 1.00 (0.43) | 1.00 (0.0.58) |
| $I/\sigma(I)$ | 16.70 (0.95) | 11.21 (1.50) | 11.85 (1.45) | 10.71 (0.89) | 7.77 (1.14) |
| Completeness (%) | 99.38 (94.80) | 99.52 (95.49) | 98.93 (89.27) | 99.67 (97.18) | 99.60 (98.65) |
| Redundancy | 12.9 (12.5) | 13.0 (12.4) | 13.1 (12.1) | 6.8 (3.9) | 10.5 (8.7) |
| **Refinement** | | | | | |
| Reflections used in refinement | 34339 (3244) | 25525 (2411) | 28595 (2528) | 41372 (3962) | 21110 (2048) |
| Reflections used for $R_{free}$ | 1720 (163) | 1280 (124) | 1430 (124) | 2070 (199) | 1057 (103) |
| $R_{work}$ | 0.17 (0.37) | 0.15 (0.28) | 0.15 (0.30) | 0.16 (0.32) | 0.16 (0.33) |
| $R_{free}$ | 0.20 (0.39) | 0.19 (0.31) | 0.19 (0.35) | 0.19 (0.35) | 0.21 (0.39) |
| CC (work) | 0.97 (0.75) | 0.97 (0.75) | 0.97 (0.81) | 0.97 (0.68) | 0.95 (0.71) |
| CC (free) | 0.96 (0.72) | 0.96 (0.78) | 0.96 (0.64) | 0.97 (0.85) | 0.96 (0.75) |
| No. of refined non-hydrogen atoms | | | | | |
| Protein | 1787 | 1793 | 1793 | 1802 | 1785 |
| Ligand(s) | 37 | 33 | 147 | 122 | 135 |
| Solvent | 117 | 216 | 234 | 237 | 209 |
| *B*-factors | | | | | |
| Protein | 34.95 | 28.57 | 24.31 | 22.80 | 29.26 |
| Lignad(s) | 54.11 | 62.26 | 35.06 | 48.48 | 46.25 |
| Solvent | 38.43 | 38.41 | 35.10 | 34.60 | 38.26 |
| Wilson *B*-factor | 25.90 | 25.39 | 22.33 | 20.17 | 26.52 |
| r.m.s.d. | | | | | |
| Bond lengths (Å) | 0.01 | 0.01 | 0.01 | 0.01 | 0.01 |
| Bond lengths (°) | 0.98 | 1.04 | 1.13 | 1.11 | 1.15 |
| Ramachandran plot | | | | | |
| Favored | 95.98 | 97.33 | 96.44 | 97.78 | 96.44 |
| Allowed | 4.02 | 2.67 | 3.56 | 2.22 | 3.56 |
| Outliers | 0.00 | 0.00 | 0.00 | 0.00 | 0.00 |
| Rotamer outliers | 1.49 | 0.00 | 0.00 | 0.00 | 0.00 |
| No. of TLS groups | 3 | 3 | 3 | 3 | 3 |

**Supplementary Table 1**: X-ray data collection and refinement statistics.

| **Enzyme** | ***Ps*Alg7A-Y223F** | ***Ps*Alg7A-Y223F** | ***Ps*Alg7C** | ***Ps*Alg7C** | ***Ps*Alg7C** |
|---|---|---|---|---|---|
| **Ligand** | **TriM** | **DiM** | **-** | **HexaM** | **PentaM** |
| **PDB ID** | 7O6H | 7NM6 | 8C3X | 8PC8 | 8PC3 |
| **Crystal condition** | 20% PEG 1500 and 0.1 M TBG buffer pH 4 | 20% PEG 1500 and 0.1 M TBG buffer pH 4 | 25% PEG3350, 0.1M Bis-Tris pH 5.5, 0.5M NaCl | 30% PEG3350, 0.1M Bis-Tris pH 5.5, 0.3M NaCl | 30% PEG3350, 0.1M Bis-Tris pH 5.5, 0.3M NaCl |
| **Protein:reservoir ratio** | 2:1 | 2:1 | 1:1 | 1:1 | 1:1 |
| **Cryoprotectant** | PEG1500 | PEG1500 | PEG400 | PEG400 | PEG400 |
| **Beamline** | BioMax | BioMax | P14 | P13 | P13 |
| **Data collection** | | | | | |
| Wavelength (Å) | 0.98 | 0.98 | 0.69 | 0.98 | 0.98 |
| Resolution (Å) | 36.17–1.66 (1.72-1.66) | 40.83–1.78 (1.89–1.78) | 33.93–0.82 (0.85–0.82) | 34.43 –1.24 (1.28–1.24) | 37.90–1.10 (1.14–1.10) |
| Space group | $P2_12_12_1$ | $P2_12_12_1$ | $P2_12_12_1$ | $P2_12_12_1$ | $P2_12_12_1$ |
| Unit cell | | | | | |
| *a, b, c* (Å) | 35.0, 80.8, 81.2 | 35.3, 80.7, 81.7 | 41.2; 59.8; 90.1 | 42.1; 59.9; 89.8 | 41.9; 59.8; 89.2 |
| Total no. of reflections | 204661 (19363) | 153775 (8545) | 2856728 (241161) | 838403 (83399) | 841521 (76052) |
| No. of unique reflections | 27861 (2706) | 22553 (1890) | 217369 (21492) | 65043 (6412) | 91405 (9008) |
| $R_{merge}$ | 0.07 (2.25) | 0.10 (0.84) | 0.05 (3.13) | 0.04 (1.76) | 0.05 (1.20) |
| $R_{meas}$ | 0.08 (2.42) | 0.11 (0.95) | 0.05 (3.27) | 0.04 (1.84) | 0.06 (1.28) |
| $R_{pim}$ | 0.03 (0.89) | 0.04 (0.44) | 0.01 (0.96) | 0.01 (0.50) | 0.02 (0.43) |
| $CC_{1/2}$ | 1.00 (0.46) | 1.00 (0.89) | 1.00 (0.29) | 1.00 (0.65) | 1.00 (0.63) |
| $I/\sigma(I)$ | 13.18 (0.91) | 10.09 (1.29) | 20.66 (0.91) | 24.63 (1.47) | 17.29 (1.43) |
| Completeness (%) | 99.85 (99.45) | 97.65 (83.52) | 99.91 (99.46) | 99.94 (100.00) | 99.93 (99.80) |
| Redundancy | 7.3 (7.4) | 6.8 (4.5) | 13.1 (11.2) | 12.9 (13.0) | 9.2 (8.4) |
| **Refinement** | | | | | |
| Reflections used in refinement | 27928 (2702) | 22547 (1890) | 217293 (21407) | 65039 (6413) | 91404 (9008) |
| Reflections used for $R_{free}$ | 1397 (135) | 1127 (94) | 1995 (198) | 1097 (108) | 1094 (108) |
| $R_{work}$ | 0.18 (0.34) | 0.16 (0.35) | 0.13 (0.30) | 0.16 (0.32) | (0.15 (0.30) |
| $R_{free}$ | 0.21 (0.33) | 0.20 (0.42) | 0.14 (0.30) | 0.17 (0.33) | 0.16 (0.29) |
| CC (work) | 0.38 (0.04) | 0.97 (0.87) | 0.97 (0.65) | 0.98 (0.77) | 0.98 (0.81) |
| CC (free) | 0.30 (0.01) | 0.89 (0.79) | 0.96 (0.58) | 0.97 (0.77) | 0.98 (0.91) |
| No. of refined non-hydrogen atoms | | | | | |
| Protein | 1775 | 1769 | 1996 | 1900 | 1961 |
| Lignad(s) | 75 | 74 | 4 | 120 | 124 |
| Solvent | 114 | 216 | 411 | 289 | 328 |
| *B*-factors | | | | | |
| Protein | 36.12 | 31.51 | 9.69 | 23.96 | 15.66 |
| Lignad(s) | 38.24 | 68.91 | 33.42 | 48.17 | 36.76 |
| Solvent | 38.59 | 39.34 | 35.33 | 36.21 | 29.35 |
| Wilson *B*-factor | 28.63 | 26.78 | 8.66 | 20.03 | 13.46 |
| r.m.s.d. | | | | | |
| Bond lengths (Å) | 0.01 | 0.01 | 0.01 | 0.01 | 0.01 |
| Bond lengths (°) | 1.02 | 1.00 | 1.06 | 1.01 | 1.10 |
| Ramachandran plot | | | | | |
| Favored | 95.54 | 95.09 | 96.49 | 96.07 | 96.93 |
| Allowed | 4.46 | 4.91 | 3.51 | 3.93 | 3.07 |
| Outliers | 0.00 | 0.00 | 0.00 | 0.00 | 0.00 |
| Rotamer outliers | 0.00 | 0.50 | 0.00 | 0.48 | 0.00 |
| No. of TLS groups | 3 | 3 | - | - | - |

**Supplementary Table 1**: X-ray data collection and refinement statistics.

| **Enzyme** | ***Ps*Alg7C** | ***Ps*Alg7C** | ***Ps*Alg7C** | ***Ps*Alg7C-Y220F** | ***Ps*Alg7C-Y220F** |
|---|---|---|---|---|---|
| **Ligand** | **TetraM** | **TriM** | **DiM** | - | **HexaM** |
| **PDB ID** | 8PCX | 8PED | 8PDT | 8C0M | 8BJO |
| **Crystal condition** | 30% PEG3350, 0.1M Bis-Tris pH 5.5, 0.3M NaCl | 30% PEG3350, 0.1M Bis-Tris pH 5.5, 0.3M NaCl | 30% PEG3350, 0.1M Bis-Tris pH 5.5, 0.2M NaCl | 2 M Ammonium sulfate; 0.1 Sodium acetate pH 4.6 | 2 M Ammonium sulfate; 0.1 Sodium acetate pH 4.6 |
| **Protein:reservoir ratio** | 1:1 | 1:1 | 1:1 | 2:1 | 2:1 |
| **Cryoprotectant** | PEG400 | PEG400 | PEG400 | PEG400 | PEG400 |
| **Beamline** | P13 | P13 | BioMax | BioMax | P14 |
| **Data collection** | | | | | |
| Wavelength (Å) | 0.98 | 0.98 | 0.80 | 0.98 | 0.77 |
| Resolution (Å) | 38.21–1.14 (1.18–1.14) | 38.78–1.10 (1.16–1.10) | 37.77–1.09 (1.16–1.09) | 20.31–1.2 (1.24– 1.20) | 19.05–1.51 (1.56–1.51) |
| Space group | $P2_12_12_1$ | $P2_12_12_1$ | $P2_12_12_1$ | *P*1 | *P*1 |
| Unit cell | | | | | |
| *a, b, c* (Å) | 42.2; 60.1: 89.8 | 42.1; 59.8; 89.3 | 41.7; 59.9; 89.0 | 34.0; 40.0; 42.6 | 33.9; 39.8; 42.6 |
| *α, β, γ* (°) | | | | 74.2; 77.1; 69.8 | 74.1; 77.9; 69.8 |
| Total no. of reflections | 1019649 (68350) | 1125679 (98256) | 659980 (61979) | 200821 (16215) | 108742 (9454) |
| No. of unique reflections | 83390 (7964) | 92064 (9095) | 92168 (8956) | 57526 (4767) | 28650 (2660) |
| $R_{merge}$ | 0.05 (1.93) | 0.06 (2.05) | 0.11 (1.7) | 0.08 (0.71) | 0.11 (0.49) |
| $R_{meas}$ | 0.05 (2.05) | 0.06 (2.15) | 0.12 (1.88) | 0.10 (0.84) | 0.12 (0.58) |
| $R_{pim}$ | 0.01 (0.68) | 0.02 (0.64) | 0.04 (0.70) | 0.05 (0.45) | 0.06 (0.30) |
| $CC_{1/2}$ | 1.00 (0.40) | 1.00 (0.45) | 1.00 (0.43) | 0.99 (0.75) | 1.00 (0.86) |
| $I/\sigma(I)$ | 20.34 (1.00) | 16.18 (1.15) | 6.89 (1.08) | 9.31 (2.10) | 8.35 (1.94) |
| Completeness (%) | 99.22 (95.91) | 99.97 (99.98) | 98.44 (96.61) | 91.28 (75.48) | 91.1 (83.7) |
| Redundancy | 12.2 (8.6) | 12.2 (10.8) | 7.2 (6.9) | 3.5 (3.4) | 3.8 (3.6) |
| **Refinement** | | | | | |
| Reflections used in refinement | 83383 (7965) | 92063 (9095) | 92159 (8956) | 57269 (4751) | 28577 (2651) |
| Reflections used for $R_{free}$ | 2100 (200) | 2096 (207) | 2100 (204) | 2887 (252) | 1458 (133) |
| $R_{work}$ | 0.12 (0.29) | 0.13 (0.28) | 0.15 (0.31) | 0.14 (0.22) | 0.17 (0.24) |
| $R_{free}$ | 0.13 (0.28) | 0.15 (0.29) | 0.19 (0.33) | 0.17 (0.25) | 0.20 (0.29) |
| CC (work) | 0.98 (0.73) | 0.98 (0.78) | 0.98 (0.75) | 0.97 (0.90) | 0.96 (0.93) |
| CC (free) | 0.98 (0.75) | 0.96 (0.76) | 0.96 (0.72) | 0.96 (0.91) | 0.96 (0.79) |
| No. of refined non-hydrogen atoms | | | | | |
| Protein | 1936 | 1847 | 1811 | 1796 | 1781 |
| Ligand(s) | 120 | 140 | 140 | 22 | 122 |
| Solvent | 373 | 346 | 353 | 339 | 228 |
| *B*-factors | | | | | |
| Protein | 19.56 | 18.24 | 16.18 | 12.54 | 16.67 |
| Ligand(s) | 31.58 | 27.28 | 32.92 | 30.30 | 26.20 |
| Solvent | 41.69 | 35.83 | 40.20 | 31.12 | 23.53 |
| Wilson *B*-factor | 15.72 | 15.34 | 13.33 | 10.18 | 13.20 |
| r.m.s.d. | | | | | |
| Bond lengths (Å) | 0.01 | 0.01 | 0.01 | 0.01 | 0.01 |
| Bond lengths (°) | 1.18 | 1.54 | 1.45 | 0.97 | 1.12 |
| Ramachandran plot | | | | | |
| Favored | 97.39 | 97.37 | 96.07 | 97.79 | 96.9 |
| Allowed | 2.61 | 2.63 | 3.93 | 2.21 | 3.10 |
| Outliers | 0.00 | 0.00 | 0.00 | 0.00 | 0.00 |
| Rotamer outliers | 1.38 | 0.50 | 0.50 | 0.00 | 0.00 |
| No. of TLS groups | - | - | - | - | 4 |

**Supplementary Table 1**: X-ray data collection and refinement statistics.

| **Enzyme** | ***Ps*Alg7C-Y220F** | ***Ps*Alg7C-Y220F** | ***Ps*Alg7C-H110N** | ***Ps*Alg7C-H110N** | ***Ps*Alg7C-H110N** |
|---|---|---|---|---|---|
| **Ligand** | **PentaM** | **DiM** | **-** | **HexaM** | **PentaM** |
| **PDB ID** | 8BXZ | 8P6O | 8RBI | 8QMJ | 8QIZ |
| **Crystal condition** | 2 M Ammonium sulfate; 0.1 Sodium acetate pH 4.6 | 2 M Ammonium sulfate; 0.1 Sodium acetate pH 4.6 | 30% PEG3350, 0.1M Bis-Tris pH 5.5, 0.2M NaCl | 30% PEG3350, 0.1M Bis-Tris pH 5.5, 0.2M NaCl | 30% PEG3350, 0.1M Bis-Tris pH 5.5, 0.2M NaCl |
| **Protein:reservoir ratio** | 2:1 | 2:1 | 2:1 | 2:1 | 2:1 |
| **Cryoprotectant** | PEG400 | PEG400 | PEG400 | PEG400 | PEG400 |
| **Beamline** | ID30B | ID30B | BioMax | BioMax | ID30B |
| **Data collection** | | | | | |
| Wavelength (Å) | 0.98 | 0.98 | 0.65 | 0.65 | 0.98 |
| Resolution (Å) | 36.62–1.2 (1.24–1.20) | 40.47–1.05 (1.09–1.05) | 0.98 | 0.98–34.49 (0.98–1.00) | 1.13–39.01 (1.13–1.16) |
| Space group | *P*1 | *P*1 | $C12_1$ | $C12_1$ | $P12_11$ |
| Unit cell | | | | | |
| *a, b, c* (Å) | 34.2; 40.0; 42.7 | 33.9; 39.9: 42.6 | 76.5; 38.5; 74.0 | 77.3; 38.8; 75.0 | 42.7; 69.4; 42.9 |
| *α, β, γ* (°) | 74.7; 78.5; 69.5 | 73.9; 77.1; 69.9 | 90.0; 116.6; 90.0 | 90.0; 116.9; 90.0 | 90.0; 114.6; 90.0 |
| Total no. of reflections | 186313 (17326) | 263305 (15734) | 755317 (46708) | 779749 (50526) | 499724 (20862) |
| No. of unique reflections | 58177 (5475) | 84331 (7347) | 110309 (6866) | 113613 (7531) | 82960 (4929) |
| $R_{merge}$ | 0.09 (0.33) | 0.11 (0.31) | 0.04 (1.40) | 0.06 (2.35) | 0.06 (1.12) |
| $R_{meas}$ | 0.11 (0.39) | 0.13 (0.39) | 0.04 (1.51) | 0.06 (2.55) | 0.07 (1.28) |
| $R_{pim}$ | 0.06 (0.22) | 0.07 (0.24) | 0.02 (0.57) | 0.02 (0.97) | 0.03 (0.60) |
| $CC_{1/2}$ | 0.99 (0.86) | 1.00 (0.96) | 1.00 (0.53) | 1.00(0.37) | 1.00 (0.62) |
| $I/\sigma(I)$ | 7.32 (4.97) | 5.97 (2.13) | 17.36 (1.29) | 13.16 (0.94) | 11.57 (1.42) |
| Completeness (%) | 92.08 (86.35) | 90.67 (78.85) | 98.64 (98.61) | 99.94 (99.87) | 96.94 (86.32) |
| Redundancy | 3.2 (3.2) | 3.1 (2.1) | 6.8 (6.8) | 6.9 (6.7) | 6.0 (4.2) |
| **Refinement** | | | | | |
| Reflections used in refinement | 58558 (5471) | 84320 (7346) | 110309 (6866) | 113613 (7531) | 82960 (4929) |
| Reflections used for $R_{free}$ | 2096 (195) | 2100 (183) | 1159 (72) | 2094 (139) | 1071 (64) |
| $R_{work}$ | 0.12 (0.12) | 0.13 (0.20) | 0.12 (0.32) | 0.12 (0.42) | 0.13 (0.34) |
| $R_{free}$ | 0.14 (0.15) | 0.15 (0.22) | 0.15 (0.36) | 0.15 (0.47) | 0.16 (0.32) |
| CC (work) | 0.97 (0.96) | 0.97 (0.93) | 0.98 (0.79) | 0.98 (0.67) | 0.98 (0.87) |
| CC (free) | 0.96 (0.93) | 0.98 (0.93) | 0.97 (0.81) | 0.97 (0.70) | 0.16 (0.32) |
| No. of refined non-hydrogen atoms | | | | | |
| Protein | 1893 | 1852 | 1881 | 1891 | 1826 |
| Ligand(s) | 103 | 49 | 8 | 49 | 122 |
| Solvent | 382 | 242 | 347 | 366 | 337 |
| *B*-factors | | | | | |
| Protein | 8.22 | 9.99 | 15.89 | 13.26 | 17.10 |
| Ligand(s) | 12.29 | 23.94 | 47.99 | 24.69 | 28.05 |
| Solvent | 24.92 | 21.04 | 34.10 | 33.58 | 37.93 |
| Wilson *B*-factor | 7.16 | 8.08 | 11.89 | 11.12 | |
| r.m.s.d. | | | | | |
| Bond lengths (Å) | 0.01 | 0.01 | 0.01 | 0.01 | 0.01 |
| Bond lengths (°) | 1.14 | 1.06 | 0.94 | 0.94 | 0.95 |
| Ramachandran plot | | | | | |
| Favored | 96.93 | 96.48 | 96.48 | 97.35 | 98.24 |
| Allowed | 3.07 | 3.52 | 3.08 | 2.65 | 1.76 |
| Outliers | 0.00 | 0.00 | 0.44 | 0.00 | 0.00 |
| Rotamer outliers | 0.00 | 0.95 | 1.42 | 1.40 | 0.98 |
| No. of TLS groups | - | - | - | - | - |

**Supplementary Table 1**: X-ray data collection and refinement statistics.

| **Enzyme** | ***Ps*Alg7C-H110N** | ***Ps*Alg7C-H110N** |
|---|---|---|
| **Ligand** | **TetraM** | **TriM** |
| **PDB ID** | 8QLI | 8R43 |
| **Crystal condition** | 30% PEG3350, 0.1M Bis-Tris pH 5.5, 0.2M NaCl | 30% PEG3350, 0.1M Bis-Tris pH 5.5, 0.2M NaCl |
| **Protein:reservoir ratio** | 2:1 | 2:1 |
| **Cryoprotectant** | PEG400 | PEG400 |
| **Beamline** | ID30B | BioMax |
| **Data collection** | | |
| Wavelength (Å) | 0.98 | 0.65 |
| Resolution (Å) | 38.67 –1.12 (1.15–1.12) | 26.83-0.87 (0.90-0.87) |
| Space group | $C12_1$ | $C12_1$ |
| Unit cell | | |
| $a, b, c$ (Å) | 77.5; 38.8; 75.2 | 77.1; 38.6; 74.6 |
| $\alpha, \beta, \gamma$ (°) | 90.0; 116.9; 90.0 | 90.0; 116.8; 90.0 |
| Total no. of reflections | 394639 (16230) | 1076500 (90602) |
| No. of unique reflections | 75472 (4449) | 159325 (15618) |
| $R_{merge}$ | 0.20 (0.48) | 0.045 (1.61) |
| $R_{meas}$ | 0.24 (0.57) | 0.048 (1.77) |
| $R_{pim}$ | 0.11 (0.29) | 0.02 (0.72) |
| $CC_{1/2}$ | 0.78 (0.70) | 1.00 (0.43) |
| $I/\sigma(I)$ | 12.98 (5.96) | 14.39 (0.85) |
| Completeness (%) | 98.33 (86.93) | 99.31 (97.65) |
| Redundancy | 5.4 (3.7) | 6.8 (5.8) |
| **Refinement** | | |
| Reflections used in refinement | 75472 (4449) | 159278 (15613) |
| Reflections used for $R_{free}$ | 2099 (124) | 2098 (206) |
| $R_{work}$ | 0.15 (0.17) | 0.12 (0.44) |
| $R_{free}$ | 0.18 (0.21) | 0.14 (0.44) |
| CC (work) | 0.91 (0.88) | 0.98 (0.74) |
| CC (free) | 0.85 (0.76) | 0.98 (0.78) |
| No. of refined non-hydrogen atoms | | |
| Protein | 1890 | 1992 |
| Lignad(s) | 49 | 61 |
| Solvent | 419 | 411 |
| *B*-factors | | |
| Protein | 9.48 | 9.95 |
| Ligand(s) | 17.52 | 12.68 |
| Solvent | 23.57 | 27.97 |
| Wilson *B*-factor | 7.65 | 9.36 |
| r.m.s.d. | | |
| Bond lengths (Å) | 0.01 | 0.01 |
| Bond lengths (°) | 1.02 | 1.05 |
| Ramachandran plot | | |
| Favored | 97.31 | 97.35 |
| Allowed | 2.69 | 2.65 |
| Outliers | 0.00 | 0.00 |
| Rotamer outliers | 0.93 | 2.17 |
| No. of TLS groups | - | - |

**Supplementary Table 2**: The sugar conformations observed in the crystal structures. Were not mentioned, the sugar moieties are in the β-conformation.

| Subsite | -3 | -2 | -1 | +1 | +2 | +3 | +4 | +5 |
|---|---|---|---|---|---|---|---|---|
| PsAlg7A (1.90 Å) | | | | | | | | |
| DP2 (1.54 Å) | | $^{4}C_{1}$ | $^{4}C_{1}$ | | | | | |
| DP3 (1.30 Å) | $^{4}C_{1}$ | $^{4}C_{1}$ | $^{4}C_{1}$ (α and β) | | | | | |
| DP4 (1.87 Å) | $^{4}C_{1}$ | $^{4}C_{1}$ | $^{4}C_{1}$ (α and β) | $\Delta$ $B_{3,O}$ | $^{4}C_{1}$ | $^{4}C_{1}$ | | |
| DP5 (1.46 Å) | $^{4}C_{1}$ | $^{4}C_{1}$ | $^{4}C_{1}$ (α and β) | $\Delta$ $B_{3,O}$ | $^{4}C_{1}$ | $^{3,O}B$ | | |
| DP6 (1.47 Å) | $^{4}C_{1}$ | $^{0}S_{2}$ | $^{4}C_{1}$ (α and β) | $\Delta$ $B_{3,O}$ | $^{1}S_{3}$ | $^{4}C_{1}$ | | |
| PsAlg7A-Y223F (1.70 Å) | | | | | | | | |
| DP2 (1.78 Å) | | $^{4}C_{1}$ | $^{4}C_{1}$ | | | | | |
| DP3 (1.66 Å) | $^{4}C_{1}$ | $^{4}C_{1}$ | $^{4}C_{1}$ (α and β) | | | | | |
| DP4 (1.81 Å) | | $^{4}C_{1}$ | $^{4}C_{1}$ | $^{2}H_{3}$ | $^{4}C_{1}$ | $^{0}S_{2}$ | | |
| DP5 (1.45 Å) | | $^{4}C_{1}$ | $^{4}C_{1}$ | $^{2}H_{3}$ | $^{4}C_{1}$ | $^{0}S_{2}$ | | |
| DP6 (1.64 Å) | $^{4}C_{1}$ | $^{4}C_{1}$ | $^{4}C_{1}$ | $^{2}H_{3}$ | $^{4}C_{1}$ | $^{4}C_{1}$ | $^{0}H_{1}$ | |
| PsPL7C (0.82 Å) | | | | | | | | |
| DP2 (1.09 Å) | | $^{4}C_{1}$ | $^{4}C_{1}$ (α) | | $^{4}C_{1}$ | $^{4}C_{1}$ | | |
| DP3 (1.10 Å) | $^{4}C_{1}$ | $^{4}C_{1}$ | $^{4}C_{1}$ (α) | | $^{4}C_{1}$ | $B_{2,5}$ | $^{1}S_{5}$ | |
| DP4 (1.14 Å) | $^{4}C_{1}$ | $^{4}C_{1}$ | $^{4}C_{1}$ (α) | | $^{4}C_{1}$ | $B_{2,5}$ | $^{1}S_{5}$ | |
| DP5 (1.10 Å) | $^{4}C_{1}$ | $^{4}C_{1}$ | $^{4}C_{1}$ (α) | | $^{4}C_{1}$ | $^{1}H_{2}$ | $^{3,0}B$ | |
| DP6 (1.24 Å) | $^{4}C_{1}$ | $^{4}C_{1}$ | $^{4}C_{1}$ (α) | | $^{4}C_{1}$ | $^{4}C_{1}$ | $^{3,O}B$ | |
| PsPL7C_Y220F (1.20 Å) | | | | | | | | |
| DP2 (1.05 Å) | | $^{4}C_{1}$ | $^{4}C_{1}$ | | $^{4}C_{1}$ (α and β) | $^{0}S_{2}$ | | |
| DP5 (1.20 Å) | | $^{4}C_{1}$ | $^{4}C_{1}$ | $^{4}C_{1}$ | $^{4}C_{1}$ | $^{3,O}B$ | | |
| DP6 (1.51 Å) | | $^{4}C_{1}$ | $^{4}C_{1}$ | $^{4}C_{1}$ | $^{4}C_{1}$ | $^{4}C_{1}$ | $^{3,O}B$ | |
| PsPL7C_H1123N (0.98 Å) | | | | | | | | |
| DP3 (0.87 Å) | $^{4}C_{1}$ | $^{4}C_{1}$ | $^{4}C_{1}$ (α) | | | | | |
| DP4 (1.00 Å) | $^{4}C_{1}$ | $^{4}C_{1}$ | $^{4}C_{1}$ | $^{1}H_{0}$ | | | | |
| DP5 (1.13 Å) | | $^{4}C_{1}$ | $^{4}C_{1}$ | $E_{2}$ | $^{4}C_{1}$ | $^{4}C_{1}$ | $^{1}S_{5}$ | |
| DP6 (0.98 Å) | | $^{4}C_{1}$ | $^{4}C_{1}$ | $E_{2}$ | $^{1}S_{5}$ | $E_{2}$ | | |

**Supplementary Table 3**: Joint X-ray and neutron data collection and refinement statistics.

| **Enzyme** | ***Ps*Alg7C** | | ***Ps*Alg7C** | |
|---|---|---|---|---|
| **Ligand** | - | | **PentaM** | |
| **PDB ID** | 8RBR | | 8RBN | |
| **Crystal condition** | 20% PEG 1500 and 0.1 M TBG buffer pH 4 | | 20% PEG 1500 and 0.1 M TBG buffer pH 4 | |
| **Protein:reservoir ratio** | 2:1 | | 2:1 | |
| **Beamline** | MaNDi | Rigaku | MaNDi | Rigaku |
| **Data collection** | | | | |
| Wavelength (Å) | 2–4 | 1.54 | 2–4 | 1.54 |
| Resolution (Å) | 13.85–2.15 (2.26–2.15) | 27.23–1.80 (1.86–1.80) | 14.56–2.15 (2.23–2.15) | 27.70–2.10 (2.16–2.10) |
| Space group | $P2_12_12_1$ | | $P2_12_12_1$ | |
| Unit cell | | | | |
| *a*, *b*, *c* (Å) | 41.58 60.97 91.31 | | 42.09 61.45 91.88 | |
| Total no. of reflections | 34916 (3145) | 278701 (24609) | 59894 (5196) | 91263 (7219) |
| No. of unique reflections | 11082 (1091) | 22174 (2168) | 12752 (1223) | 14513 (1147) |
| $R_{merge}$ | 0.19 (0.23) | 0.13 (0.44) | 0.22 (0.27) | 0.13 (0.66) |
| $R_{meas}$ | 0.22 (0.27) | 0.13 (0.46) | 0.24 (0.31) | 0.14 (0.72) |
| $R_{pim}$ | 0.11 (0.14) | 0.037 (0.14) | 0.10 (0.13) | 0.06 (0.29) |
| $CC_{1/2}$ | 0.90 (0.45) | 0.98 (0.94) | 0.94 (0.33) | 1.00 (0.80) |
| $I/\sigma(I)$ | 5.29 (3.48) | 20.70 (6.08) | 8.18 (3.31) | 12.4 (4.0) |
| Completeness (%) | 84.03 (85.10) | 99.92 (100.00) | 94.61 (92.16) | 100.0 (100.0) |
| Redundancy | 3.2 (2.9) | 12.6 (11.4) | 4.7 (4.2) | |
| **Refinement** | | | | |
| Reflections used in refinement | 11082 (705) | | 12771 (1121) | |
| Reflections used for $R_{free}$ | 549 (38) | | 633 (53) | |
| $R_{work}$ | 0.32 (0.37) | | 0.35 (0.39) | |
| $R_{free}$ | 0.32 (0.33) | | 0.36 (0.45) | |
| CC (work) | 0.76 (0.24) | | 0.72 (0.32) | |
| CC (free) | 0.84 (0.53) | | 0.74 (0.19) | |
| No. of refined non-hydrogen atoms | | | | |
| Protein | 2040 | | 1774 | |
| Ligand(s) | 0 | | 110 | |
| Solvent | 597 | | 372 | |
| *B*-factors | | | | |
| Protein | 14.43 | | 15.09 | |
| Ligand(s) | - | | 54.70 | |
| Solvent | 30.28 | | 26.13 | |
| r.m.s.d. | | | | |
| Bond lengths (Å) | 0.11 | | 0.02 | |
| Bond lengths (°) | 1.18 | | 1.71 | |
| Ramachandran plot | | | | |
| Favored | 97.82 | | 96.94 | |
| Allowed | 2.18 | | 3.06 | |
| Outliers | 0.00 | | 0.00 | |
| Rotamer outliers | 0.00 | | 1.03 | |
| No. of TLS groups | - | | - | |

**Supplementary Table 4.** List of the primers used for site directed mutagenesis. Mutated codons are underlined, f indicates forward primers and r, reverse primers.

| Primer | Sequence |
|---|---|
| PsAlg7A Y223F f | 5'-C TTC AAG GCT GGT GCT TTT AAC AAT AAC CCA ACT GAT ACT TC-3' |
| PsAlg7A Y223F r | 5'-AGC ACC AGC CTT GAA GTA ACA AGT AGA ACC ATC GAA TTG-3' |
| PsAlg7C H124N f | 5'-CT ATT ATG CAA ATT AAC TCT GGA GAG GCT C-3' |
| PsAlg7C H124 r | 5'-AAT TTG CAT AAT AGT AAC CTC TTC AAC AC-3' |
| PsAlg7C Y220F f | 5'-TTC AAG GCT GGT GCT TTT AAC AAT AAC CCT ACT TCT GAG-3' |
| PsAlg7C Y220 r | 5'-AGC ACC AGC CTT GAA ATA ACA AGA ATC AGT ACC ATC C-3' |
